# Grand canonical Brownian dynamics simulations of adsorption and self-assembly of SAS-6 rings on a surface

**DOI:** 10.1101/2022.11.19.517069

**Authors:** Santiago Gomez Melo, Dennis Wörthmüller, Pierre Gönczy, Niccolo Banterle, Ulrich S. Schwarz

## Abstract

The protein SAS-6 forms dimers, which then self-assemble into rings that are critical for the nine-fold symmetry of the centriole organelle. It has recently been shown experimentally that the self-assembly of SAS-6 rings is strongly facilitated on a surface, shifting the reaction equilibrium by four orders of magnitude compared to the bulk. Moreover, a fraction of non-canonical symmetries (i.e., different from nine) was observed. In order to understand which aspects of the system are relevant to ensure efficient self-assembly and selection of the nine-fold symmetry, we have performed Brownian dynamics computer simulation with patchy particles and then compared our results with experimental ones. Adsorption onto the surface was simulated by a Grand Canonical Monte Carlo procedure and Random Sequential Adsorption kinetics. Furthermore, self-assembly was described by Langevin equations with hydrodynamic mobility matrices. We find that as long as the interaction energies are weak, the assembly kinetics can be described well by the coagulation-fragmentation equations in the reaction-limited approximation. By contrast, larger interaction energies lead to kinetic trapping and diffusion-limited assembly. We find that selection of nine-fold symmetry requires a small value for the angular interaction range. These predictions are confirmed by the experimentally observed reaction constant and angle fluctuations. Overall, our simulations suggest that the SAS-6 system works at the crossover between a relatively weak binding energy that avoids kinetic trapping and a small angular range that favors the nine-fold symmetry.

## I. INTRODUCTION

Proteins are the working horses of biological systems and their assembly into supramolecular complexes lies at the heart of nature’s astonishing ability to build structures with specific functions^1,2^. Well-known examples for such functional complexes include the cytoskeleton made of actin, microtubules and intermediate filaments, as well as flagella and cilia, clathrin cages, nuclear pore complexes and viral capsids. For all the above examples, corresponding mathematical and computational models have been developed^3–5^. These models have revealed that assembly of large protein complexes is a challenging task due to conflicting requirements. On the one hand, the function of these complexes usually dictates a desired target structure, which from a physical point of view has to be stabilized by a large gain in free energy. On the other hand, it has been shown that fast assembly driven by large gains in free energies typically leads to kinetic trapping and malformed structures, thus rendering the entire process very inefficient^6–8^. In the cellular context, extrinsic mechanisms of target stabilization might exist, such as binding of the target structure to other proteins or post-translational modifications. Yet many important protein self-assembly reactions have been successfully reconstituted *in vitro* with minimal components, demonstrating that intrinsic mechanisms can be sufficient to ensure that assembly is both efficient and specific^9^.

Here we address this central aspect of protein assembly for such a system that has been reconstituted *in vitro*, namely assembly of Spindle Assembly Abnormal Protein 6 (SAS-6) into rings. SAS-6 is critical for the formation of centrioles, which are cylindrical nine-fold symmetric microtubule-based organelles at the core of centrosomes, the main microtubule organising centers (MTOCs) of animal cells (Fig. 1a). After cell division, each centrosome contains a centriole pair. At the G1/S-transition of the cell cycle, each of the resident centrioles seeds the formation of a nascent centriole orthogonally from a surface at its proximal end (Fig. 1b). In late G2, the two pairs migrate to the opposite side of the nucleus, thereafter forming the poles of the mitotic spindle^10^. Given its role in MTOC formation, centrioles are crucial for many essential cellular processes, including cell polarisation, division and motility^11^. As might be expected from such important cellular functions, centriole number misregulation and structural aberrations have been linked to numrous pathological conditions, including microcephaly, ciliopathies and cancer^12^.

**FIG. 1:**
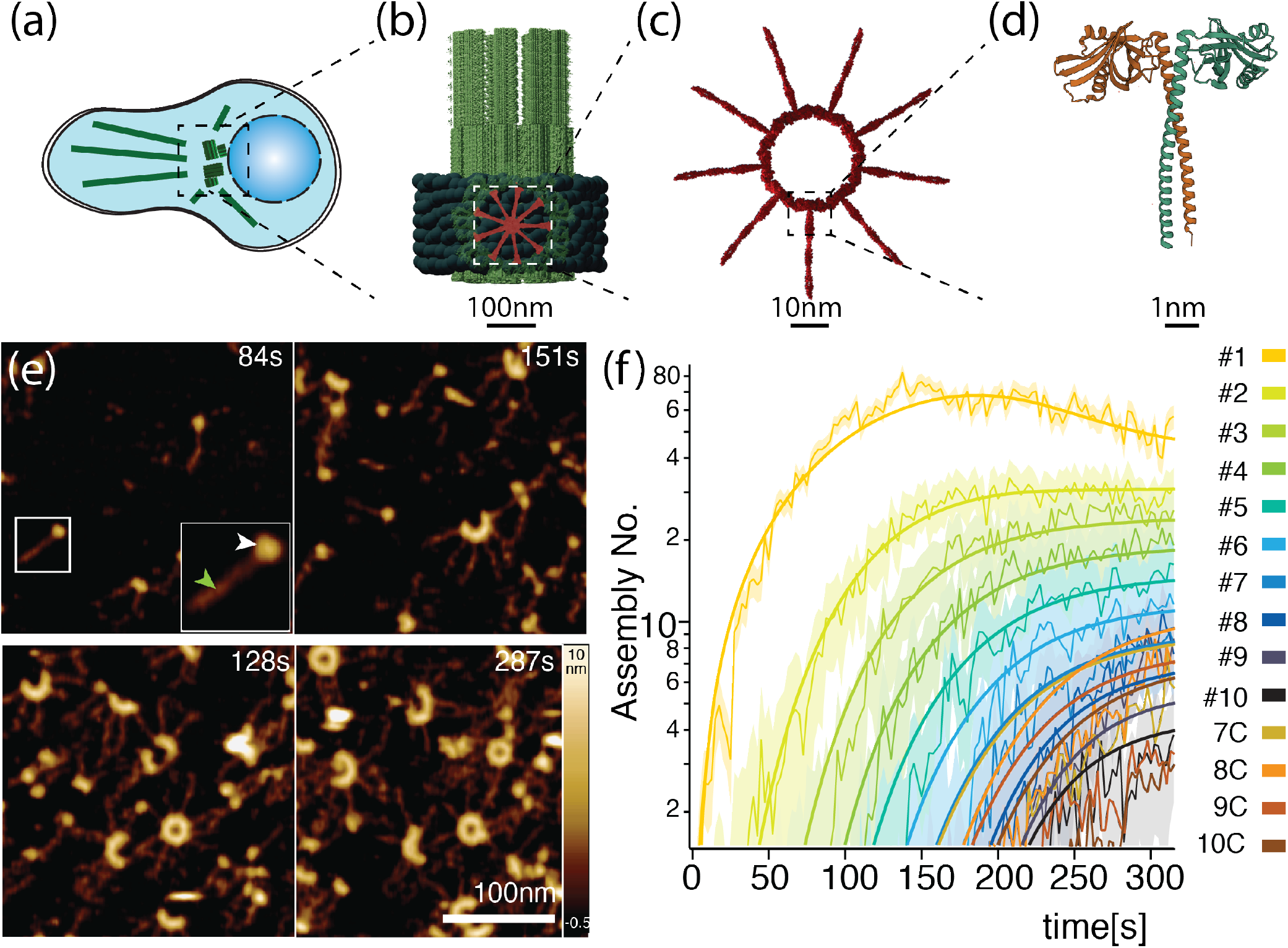
(a) Centrosomes are microtubule-organizing centers in animal cells that consist of centrioles. (b) A nascent centriole grows orthogonally from the surface of a resident centriole. (c) The ninefold symmetry of the centriole derives from a nonameric ring of SAS-6 dimers. (d) Each dimer has a coiled-coil domain responsible for homo-dimerization and a N-terminal domain responsible for higher order oligomerization. (e) Snapshot of SAS-6 assembly on a surface as observed by PORT-HS-AFM experiments. Note that for the subsequent image processing of these experiments, one must use the SAS-6 variant with full spokes, whereas the simulations hereafter are conducted with shorter spokes. Arrowheads in the inset mark the head (white) and the spoke (green) of one dimer, respectivley. (f) Time evolution of the concentrations of the differently sized SAS-6 oligomers during ring assembly. First, SAS-6 dimers are adsorbed onto the surface and then assemble into oligomers up to size ten, including closed rings from sizes seven to ten. (e) and (f) reproduced under the Creative Commons Attribution 4.0 International License from^13^.

Dimers of SAS-6 are the building blocks of the first structure present in nascent centrioles, namely the so-called centriolar cartwheel, a nine-fold symmetric element formed of a stack of protein rings and peripheral elements connecting it to the microtubule wall^14^. The cartwheel is critical for centriole formation and is instrumental in establishing its characteristic nine-fold symmetry. The presence of SAS-6 is necessary for cartwheel formation; SAS-6 proteins can assemble *in vitro* into nine-fold symmetric rings^15,16^, composed of nine dimers, exactly as in the cartwheel (Fig. 1c). Centrioles of *C. reinhardtii* bearing mutations in CrSAS-6 have symmetries deviating from the canonical nine-fold, demonstrating the fundamental role of this protein in contributing to dictate the structure of the organelle^15,17^.

At the structural level, SAS-6 is composed of a globular N-terminal domain and a coiled-coil (CC) domain, followed by a variable and potentially unstructured C-terminal moiety (Fig. 1d). The protein dimerizes via strong interactions among the CC-domains, forming a homodimer with a globular N-terminal head and a CC-domain that yields a spoke extending away from the head domain. The dimers are then able to form larger complexes, including nine-fold symmetrical ring, by means of weaker yet highly anisotropic interactions between the N-terminal domains^15,16,18^.

The dynamics of SAS-6 ring polymerisation has been visualised and characterised with Photothermal Off-Resonance Tapping High Speed Atomic Force Microscopy (PORT-HS-AFM) on mica surfaces. This allowed for the visualisation of single molecule dynamics with minimally invasive forces, whilst still retaining high spatial (in the nm range) and temporal (in the second range) resolution^13,19^. Combining the PORT-HS-AFM approach with quantitative image processing, reaction kinetics and MD-simulations enabled a detailed analysis of the structural and kinetic mechanisms of SAS-6 ring assembly^13^. Image processing of such data sets is faciliated by the presence of the long spokes of SAS-6, which can be used to assign an oligomerization state to the growing higher order oligomers (Fig. 1e). Plotting the concentrations of the different intermediates gives a full kinetic time course of the assembly reaction (Fig. 1f). These studies showed that SAS-6 self-assembly occurs by first adsorbing dimers onto the surface; these dimers then assemble into higher order oligomers (up to ten), including closed rings of sizes seven to ten.

It was also shown that the presence of the surface has a catalytic effect, shifting the equilibrium of the reaction by four orders of magnitude^13^. This means that association is greatly facilitated on a surface compared to in solution, an effect that results mainly from the fact that proteins have a higher encounter probability in two versus three dimensions. Another important aspect is the effect of the surface on the structure of the SAS-6 dimers. MD-simulations showed that SAS-6 oligomers in solution tend to form a shallow helix, but the surface forces the complex into a planar structure and thus makes ring closure possible. Together, these factors help explain why in a cellular context the nascent centriole is assembled solely on the surface of the existing centrosome and not in the cytosol. These results also suggest that SAS-6 rings are prestressed, which might be important to mechanically stabilize the centriole. Finally, it should be noted that the helical structure might contribute to break the rotational symmetry of the centriole.

In order to demonstrate the structural role of the surface in suppressing formation of a helix and stabilizing rings, allatom and coarse-grained molecular dynamics (MD) computer simulations have been used^13^. However, this approach cannot access the length and time scales required to simulate the assembly of entire rings. A standard approach to this challenge is the use of Brownian Dynamics (BD) to coarse-grain all fast degrees of freedom such as collisions with the solvent particles, which then effectively go into noise terms for the slow degrees of freedom,. In addition, the protein structure itself is coarse-grained via suitable models such as patchy particles^3,20^. However, even BD might be too computationally costly, in particular when modelling assembly of protein complexes with many intermediates. One then typically resorts to reaction kinetics (RK), which describe the time evolution of macroscopic concentrations via ordinary differential equations rather than single particles interacting in time and space. Like for coarse-graining from MD to BD, coarse-graining from BD to RK requires that one asks under which conditions this procedure is justified and under which condition it should be avoided. This important methodological question is tightly connected with the biological circumstances under which protein self-assembly can be efficient due to reversibility, because only in this case does one expect RK to work well.

Here we address these important questions in the context of SAS-6 ring assembly on a surface as shown in Fig. 1. In this case, the RK is described by the coagulation-fragmentation (CF) process, which has been extensively investigated in applied mathematics and is widely used for reversible assembly processes in chemistry and physics^21,22^. Adapted to SAS-6 ring assembly with up to ten-rings, which can be either open or closed, it was possible to fit the solutions of the CF differential equations to the PORT-HS-AFM data^13^. This was achieved with the reaction-limited version of the CF-equations, which focuses on the two model parameters *k*_+_ and *k*_−_, independently of the sizes of the reacting oligomers. However, despite its success in describing the experimentally observed kinetics, the RK-approach is in principle limited because it does not include the spatial domain. In particular, it does not account for the exact nature of adsorption from solution and for the roles of translational and rotational diffusion on the surface. Thus it is not clear what the limits are of the RK-approach in this case.

In order to test the validity of the RK-model and to assess the role of the spatial degrees of freedom, one has to turn to computer simulations in the spatial domain, for which BD is most appropriate given the large size of the system. BD-simulations of patchy particles have been used before to investigate ring formation in solution^23,24^ and have also been applied to the case of SAS-6 rings^25^. However, a major limitation of these previous BD-simulations is the absence of ring size variability^25^. Rings such as decamers or octamers, which deviate from the canonical ninefold symmetry, have been observed in the PORT-HS-AFM experiments^13^. Earlier work on ring formation with patchy particles did not allow for such structures and was only concerned with the stochastic formation of the desired nine-fold symmetrical target structures^25,26^. In order to allow for ring size variability, here we opt for a force-based approach previously used for viral capsid studies^6^, in which binding occurs via an attractive force resulting from an anisotropic interaction potential between the binding sites of the protein. This enables assembly of malformed structures (i.e., other than nine-fold) and allows for the evaluation of the impact of microscopic parameters, such as the strength of the interaction or the extent of its anisotropy, on the relative population of these oligomers.

In order to analyze ring formation on surfaces using the BD-approach, one not only has to formulate the stochastic equations in two dimensions, but also to include a theoretical treatment of dimer adsorption from solution. In order to achieve quantitative understanding of the SAS-6 assembly process on a surface with adsorption from the bulk, we have implemented a grand canonical BD-procedure for suitably coarse grained interacting SAS-6 dimers. We decided to model a truncated variant of the SAS-6 protein containing only six heptad repeats of the coiled-coil domain, for which the crystal structure has been solved^15^ and a coarsegrained computer model has been developed previously^25^. This choice allows us also to focus on the effect of dimer-dimer binding and to avoid the complications arising from longer spokes. The shape of the protein is approximated with a finite collection of spheres, whereas the rapid degrees of freedom, such as the collisions with the solvent particles, are modelled as stochastic noise^3^. The BD routine is coupled with a Grand Canonical Monte Carlo (GCMC) algorithm that accounts for particle exchange with a reservoir. The reservoir models a bulk solution which is present both in the PORT-HS-AFM experiments and in a cellular context. The resulting implementation monitors the time evolution of differently sized complexes and can be compared both to experimental and RK-results. In particular, this approach allows us to identify the limits of the RK-approach, that is the CF-equations in the reaction-limited form, and to investigate if and how the SAS-6 system can strike the balance between reversible assembly and target selection.

## II. METHODS

### A. Langevin equations and friction matrices

We simulate a two-dimensional fluid contained in a square box of length *L* which lies in the *xy* plane (Fig. 2a). Proteins in the fluid are able to move and interact among themselves. In addition, the fluid is in contact with an infinitely large protein reservoir with particle density ρ so that particle exchange via adsorption and desorption processes is possible. The motion of the particles in the fluid is described by overdamped Langevin dynamics, whereas the adsorption process is simulated via a Grand Canonical Monte Carlo routine.

**FIG. 2:**
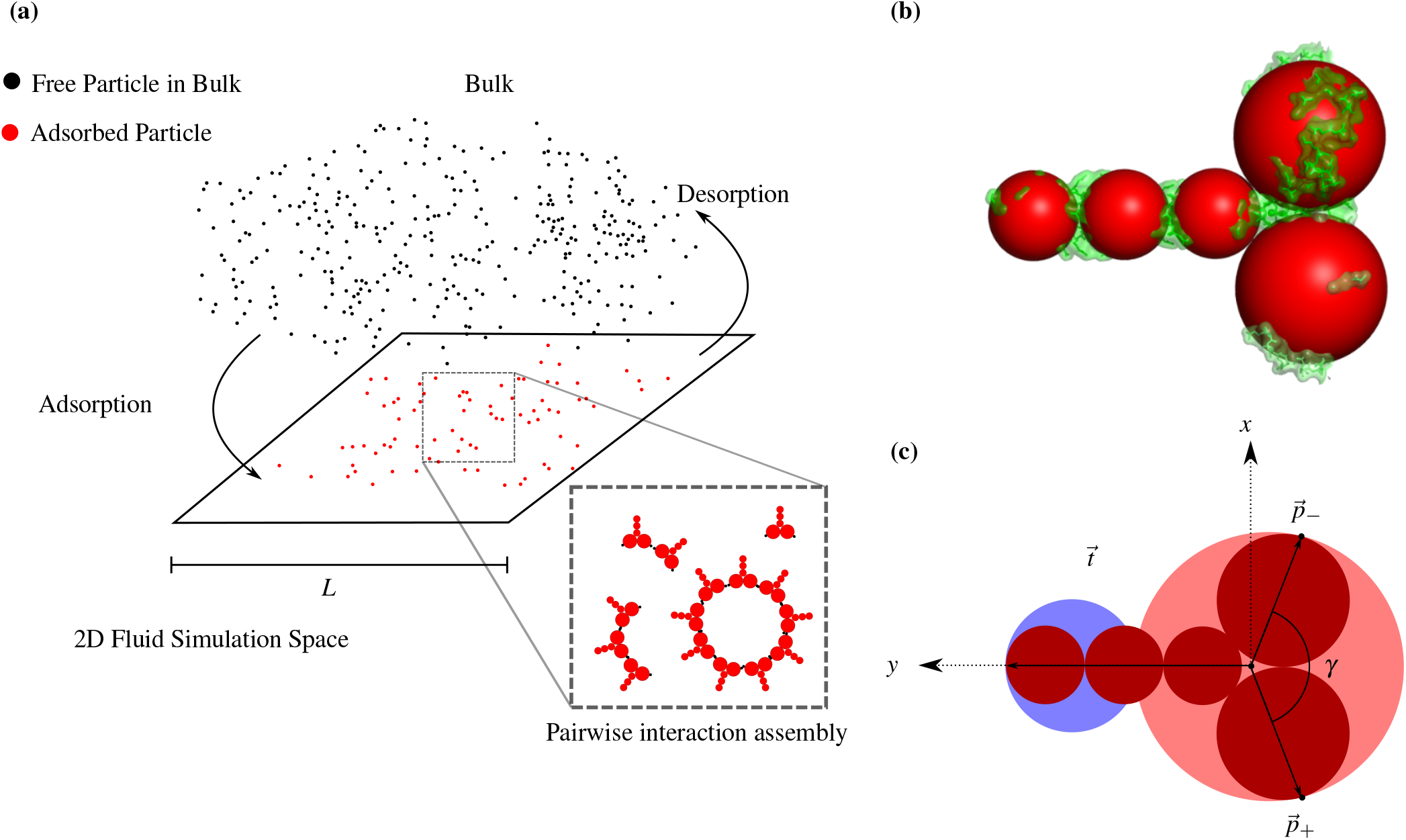
(a) The simulated system consists of a two-dimensionalfluid contained in a square box of size *L* with interacting proteins. Thefluid is in contact with a particle reservoir which allows for particle exchange. (b) Coarse-grained model of a SAS-6 homodimer with six heptad repeats (short spokes) proposed by Klein^25^, superimposed with its X-ray structure. This model is used to calculate the diffusion matrix. (c) A second coarse-grained model with two rather than six beads is used to simulate assembly. In the particle fixed frame, the *y*-axis is aligned with the CC-domain. The bigger (red) sphere models the head domain and the smaller (blue) sphere models the coiled-coil domain. Two patches are placed symmetrically at positions 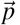± to act as binding sites, and the angle *γ* = 140° is set so that perfect alignment of two patch vectors corresponds to the 40° angle of a regular nonagon.

The trajectory of each protein in the two dimensional fluid is resolved according to the Langevin equation of motion in the overdamped limit^23,27^

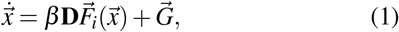

Where 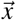 is the coordinate vector of the particle, 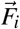 is the generalized total force arising from protein-protein interactions, 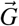 is Gaussian noise accounting for rapid degrees of freedom such as collision with solvent particles, **D** is the diffusion matrix and β = 1/*k*_*B*_*T*. In the two dimensional case the proteins possess three degrees of freedom: two for their position in the simulation domain, specified by 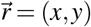, and one for the orientation angle *α*. Hence, all vector quantities in Eq. 1 are three dimensional, and 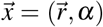. Similarly, 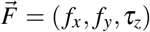 contains in-plane translational forces and an out of plane torque. The diffusion matrix is 3 × 3. If there are *N* particles on the surface, the algorithm solves 3*N* coupled stochastic differential equations.

The moments of the Gaussian noise are specified by the diffusion matrix via the expressions^23,27^

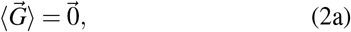

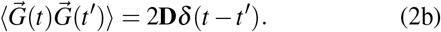

One therefore requires knowledge of the diffusion matrix and the interaction forces in order to solve Eq. 1.

The diffusion matrix is naturally defined in the co-moving frame attached to the protein, also known as the particle fixed frame (PFF). Although the interaction with the surface changes the diffusion matrix of molecules in the bulk, we still expect that essential features like the relative importance of translation versus rotation to be similar. The diffusion matrix is therefore estimated by means of the bead model proposed by de la Torre and Carrasco^28,29^ combined with the modified Kirkwood-Riseman treatment as implemented by de la Torre and Bloomfield^30^. This model allows for the computation of the diffusion matrix in the Stokes hydrodynamic regime for a rigid body composed of *N* co-moving spheres. De la Torre implemented these methods in the HYDRO++ program^31^ which was here used to compute the diffusion matrix of SAS-6. In order to do so, the molecule was coarse-grained with five non-overlapping spheres (Fig. 2b) following earlier work by Klein^25^.

The HYDRO++ program yields 36 components in four matrices of dimension 3 × 3 because it considers a three dimensional molecule in a three dimensional fluid. Since the algorithm restricts itself to two dimensions, only nine components out of the 36 entries, corresponding to the entries of **D**, are relevant. Moreover, the off-diagonal elements are at least two orders of magnitude smaller than the diagonal elements, so they can be neglected. The final diffusion matrix in the PFF is therefore diagonal, given by

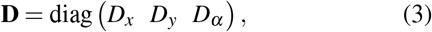

where *D*_*x*_ = 6.621 ×10^−7^cm^2^ s^−1^, *D*_*y*_ = 6.97 × 10^−7^cm^2^ s^−1^ and *D*_*α*_ = 2.04 × 10^6^s^−1^. The full matrix as predicted by HYDRO++ is reported in the supplementary material.

Next, the interaction forces are explicitly introduced. The full coarse grained model in Fig. 2b would be computationally expensive since one had to check all pairwise interactions for 5*N* spheres. Therefore, for the calculation of interaction forces, SAS-6 is further reduced to two contacting spheres with two patches as binding sites (Fig. 2c). The two spheres correspond to the head and the coiled-coil, respectively, and have corresponding radii of *R*_*b*_ = 8.5nm and *R*_*t*_ = 3.5nm. A free body diagram of two interacting proteins is shown in Fig. 3.

**FIG. 3:**
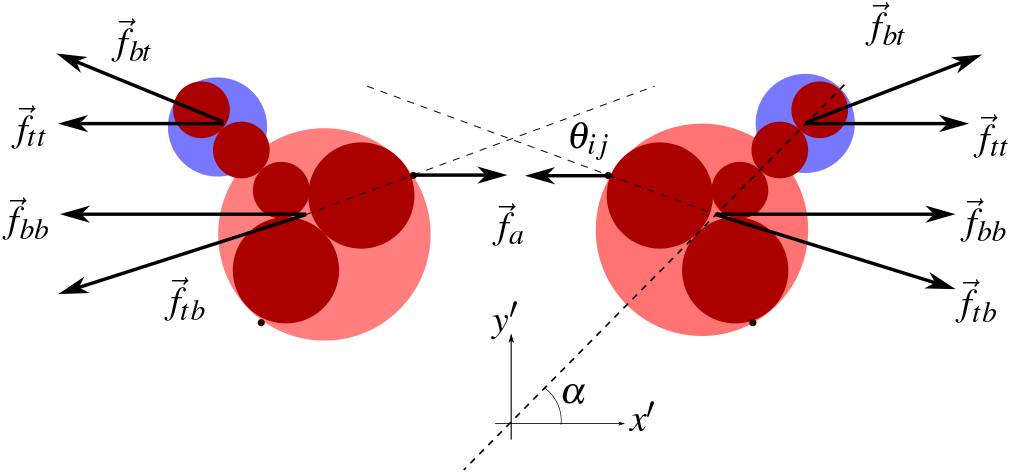
Interactions between a pair of SAS-6 dimers in the lab frame. They exhibit four types of repulsive interactions: between heads (*bb*), coiled-coil (*tt*) and heads and coiled-coil (*tb* and *bt*). In addition, the binding sites lead to patch alignment with the canonical nine-fold symmetry being most favorable. The particle orientation is measured with respect to the lab frame via an angle *α* which defines the orthogonal transformation to the particle fixed frame.

Each pair of proteins may experience five types of interaction forces: four arising from repulsion between heads (*bb*), coiled-coil (*tt*) and head-coiled-coil (*tb* and *bt*), and an attractive one between patches, which are labelled as + or −. For the repulsive interactions, the explicit form of the potential is given by a Weeks-Chandler-Andersen function, motivated by studies on viral capsid assembly^6^

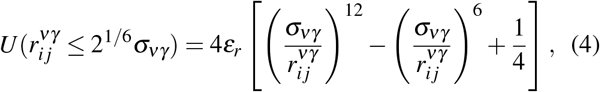

and 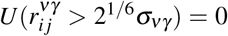. In Eq. 4, 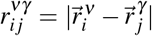, is the distance between the spheres *i* and *j*, and ν, *γ* ∈ {*b, t*} specify if the sphere corresponds to a coiled-coil or a head. Additionally, ε_*r*_ defines the strength of the interaction and σ_ν*γ*_ the length scale of the interaction. The potential is truncated at its minimum, namely at a cutoff distance *c*_*νγ*_ = 2^1/6^σ_*νγ*_, so that it does not has any attractive part, but smoothly becomes 0 for larger distances. The total repulsive force on sphere ν of particle *i* then reads

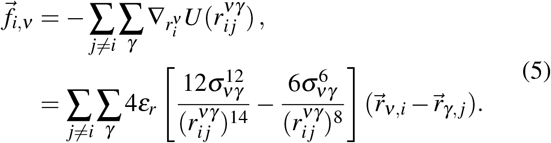

The total repulsive force on a protein is then the sum of the forces in the head and coiled-coil. Furthermore, the forces on a tail will exert a torque given by

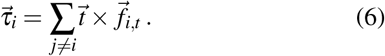

On the other hand, the attractive interaction between patches *U*_*a*_ is chosen to be a product of a radial function *u*(*r*_*i j*_) and an alignment switch function *s*(*θ*_*i j*_), again motivated by viral capsid assembly^6^

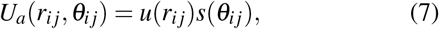

where *r*_*i j*_ is the distance between patches. The radial dependence is given by shifted Lennard Jones potential truncated at a certain cutoff *r*_*c*_ with binding energy *ε*_*a*_ and the same length scale as that of body-body repulsive interaction σ_*bb*_

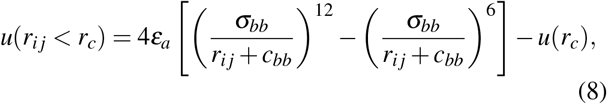

where *c*_*bb*_ = 2^1/6^σ_*bb*_ is the repulsive cutoff for body-body interaction and *r*_*c*_ = 2.5σ_*bb*_− *c*_*bb*_. Likewise, *u*(*r*_*ij*_≥ *r*_*c*_) = 0. The constant term *u*(*r*_*c*_) ensures continuity at *r* = *r*_*c*_. Moreover, the angular function is written in terms of a truncated cosine,

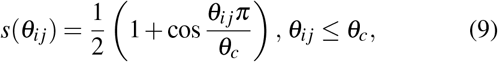

and *s*(*θ*_*ij*_ ≥ *θ*_*c*_) = 0 for a cutoff angle *θ*_*c*_. This function decays monotonously from 1 to 0 between for *θ* ≤ *θ*_*c*_ and then becomes 0 smoothly everywhere else. Consequently, *s*(*θ*) acts as a selector that restricts possible binding partners to those which deviate from perfect alignment at most by an angle *θ*_*c*_, thus accounting for bonding anisotropy. Graphs for the interaction potentials are provided in the supplementary material.

When computing the attractive force field, it is important to distinguish between the + and − patches; in the PORT-HS-AFM-experiments, one only observes bonding between the patches of different labels, leading to ring geometries. Bonding between patches of equal labels would lead to zig-zag configurations which are observed to be only short-lived experimentally, so they are not taken into account in our simulations.

The force on the + patch of the *i*-th particle is given by

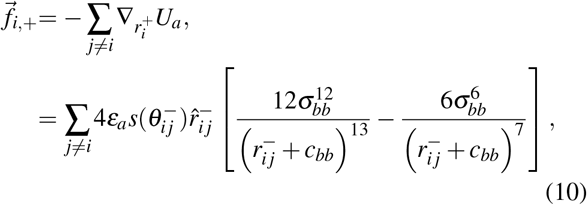

where 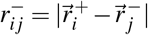 is the distance to the − patch of the *j*-th particle, 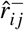 is the unit vector connecting these patches and 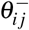 is the bond angle, which is calculated as

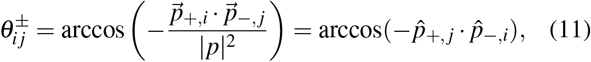

where 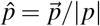 denotes unitary vectors. The − superscripts on the right hand side of Eq. 10 are chosen to remind one that only interactions with patches of that label have to be computed for a + patch. The force for the − patch is given by the same equation, but with flipped labels.

Attractive interactions will also exert a torque on the particles, given by

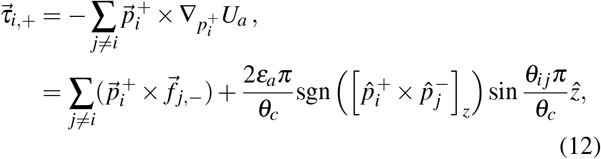

for a + patch, where sgn is the sign function.

Now that the explicit forms of the interaction forces and diffusion matrix are known, the Langevin equations of motion may be integrated. However, the diffusion matrix in Eq. 3 is defined in the PFF. This diagonal form of the matrix is advantageous, as it allows for efficient vectorization in the code and the factoring of the distribution of the components of 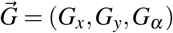 into a product of single variable distributions. Nonetheless, the goal of the routine is to calculate protein positions and orientations in a lab frame at rest, whereas the PFF rotates with each individual protein. Therefore, a rotation operator in the *xy* plane **R**(*α*) is needed to transform between these frames, where *α* quantifies the particle orientation in the lab frame as shown in Fig. 3. This rotation only transforms the first two in-plane components. The Langevin equation for the velocity in the lab frame denoted by 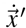 is given by

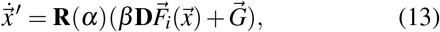

where the components of all unprimed quantities are measured in the PFF. Eq. 13 is solved numerically using a forward Euler integrator with periodic boundary conditions and minimum distance convention. The discretization of the differential equation results in a finite time step Δ*t*. This discrete time step impacts the second moment of the probability density of 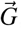, which now reads

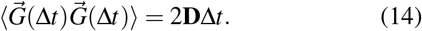

A time step of 25ps is chosen because it gives a good compromise between the diffusion and interaction force length scales while allowing for the observation of the system evolution in acceptable computational time. Thus far, the simulation of SAS-6 self assembly is possible for a constant, non-zero number of dimers. Next, we couple the simulation with a protein reservoir with the aid of a Grand Canonical Monte Carlo routine.

### B. Grand canonical Monte Carlo scheme

To simulate the adsorption process on a surface, a Grand Canonical Monte Carlo algorithm is implemented. The algorithm is based on the equilibrium marginal probability distribution 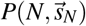 of finding *N* adsorbed particles at rescaled positions 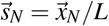, which is given by the Boltzmann factor in the Grand Canonical ensemble

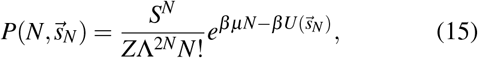

where *μ* is the chemical potential, 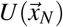 is the total potential energy of the adsorbed particles, *Z* is the normalizing partition function, *S* = *L*^2^ is the area of the square and Λ is the thermal de Broglie wavelength for a particle of mass *m*,

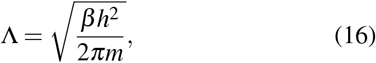

where *h* is Planck’s constant. The Λ^2*N*^ factor in the denominator of Eq. 15 comes from tracing out the momenta of all the particles from the full Boltzmann distribution. Similarly, the *S*^*N*^ factor appears when rescaling the positions as a fraction of the box length *L*, so the probability density must be multiplied by the Jacobian of the transformation in order to preserve the measure of the probability space.

This equilibrium distribution allows for the computation of transition amplitudes *k*(*N, N* + 1) from a state with *N* particles to another with *N* + 1 adsorbed particles. In order to reach the correct probability density given by Eq. 15, these transitions must obey detailed balance^32^, so

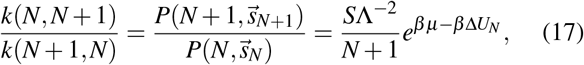

where 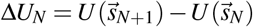 is the potential energy difference arising from the additional particle. A similar expression may be derived for a single particle desorption by replacing *N* → *N* − 1. Once this ratio is specified, a Metropolis Hastings type algorithm is implemented^32^, so the transition probabilities are

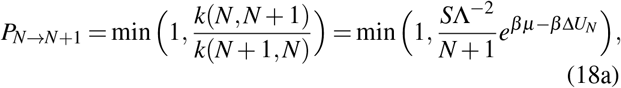

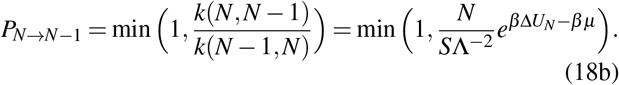

In an adsorption event, the particle is created at a random position, and in the case of desorption an existing particle is selected at random. The algorithm may therefore be summarized in the following steps^32^:

**1:** Choose a random position where particle creation is proposed, or select a random adsorbed particle whose annihilation is attempted.

**2:** Calculate the interaction energy difference of this event Δ*U*_*N*_.

**3:** Calculate acceptance probability *P*_*a*_ according to Eq. 18a for adsorption and Eq. 18b for desorption.

**4:** Draw a number *z* from the uniform distribution *U* (0, 1) and accept the event if *z* < *P*_*a*_.

In order to implement this algorithm, two quantities must be specified: the chemical potential *μ* and the energy Δ*U*_*N*_. The chemical potential is approximated to that of an ideal gas, which is given by the Sackur Tetrode equation,

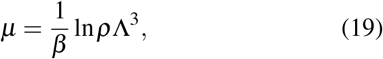

where ρ is the particle number concentration of the reservoir. This approximation is adequate for dilute solutions of weakly interacting particles, such as SAS-6. Nevertheless, other models for chemical potential are readily available to simulate adsorption from more realistic bulk solutions.

The potential energy difference is calculated as the sum of the pairwise repulsive interactions of all the particles plus an additional constant *V*_0_ term accounting for attractive interactions with the surface,

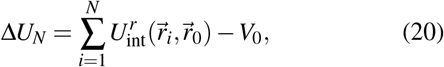

where 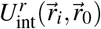 is the pairwise repulsive interaction between particles *i* and the test particle to be created or destroyed, which is labelled as the 0 particle without loss of generality. This interaction potential is the sum of the previously discussed interactions on coiled-coil and head, as given by Eq. 4. The additional *V*_0_ term arising from interactions with the surface is introduced so that adsorption becomes energetically favourable and dimers in the bulk are attracted to the surface. In the case of the SAS-6 experiments on a surface, this term models mainly the electrostatic interaction between the negatively charged mica and the dimers suspended in solution. With the explicit form of the potential, now the algorithm may simulate fluctuating particle number and adsorption process in addition to the in-plane protein motion.

### C. Adsorption kinetics and RSA

So far no reference to time has been made in the modelling of adsorption. This is understandable since detailed balance only ensures that the right probability distribution is sampled at equilibrium; how fast the system equilibrates is quite a different matter. Defining an adsorption-related time scale is important because it establishes how fast adsorption triggers the assembly on the surface. Therefore, a second time step is introduced in order to gauge the rate of adsorption onto the surface. This time step, referred to as GCMC time step or Δ*t*_GCMC_, is defined as an integer multiple of the original time step for Brownian dynamics

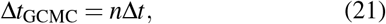

so that both an adsorption and a desorption event is computed every *n* time steps. This allows the user to define the number of GCMC attempts per unit time. A lower *n* means that more GCMC events are attempted in a given time interval, thus resulting in a higher adsorption.

In order to enable comparison with macroscopic rate equations, a rate model describing adsorption is required. The most suitable model for the algorithm is the *Random Sequential Adsorption* or RSA in the limit of low coverage, which is given by a second order polynomial^33^,

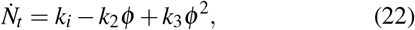

where *N*_*t*_ is the total number of adsorbed particles and 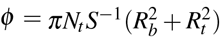 is the surface coverage. This explicit form was derived by Schaaf and Talbot^33^, who analytically calculated the coefficients *k*_2_ and *k*_3_ for the case of hard spheres. In our case, *k*_2_ and *k*_3_ are treated as free parameters to be fitted to the simulation results. *k*_*i*_ may be approximated as *k*_*i*_ ≈ 1/Δ*t*_GCMC_ as long as the initial adsorption probability 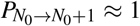, which is the case of all simulations. The explicit form of the adsorption kinetics enables the comparison between the algorithm and macroscopic rate models, which are introduced next.

**D. Coagulation-fragmentation equations for rings**

The microscopic simulation is to be compared with macroscopic reaction kinetics (RK). The dynamics of protein self-assembly, such as that of SAS-6, are often described by the coagulation-fragmentation (CF) equations^21,22^. Here, we not only incorporate ring formation^13^, but also add an additional source term *g*(*N*_*t*_) to incorporate adsorption from a reservoir, to arrive at

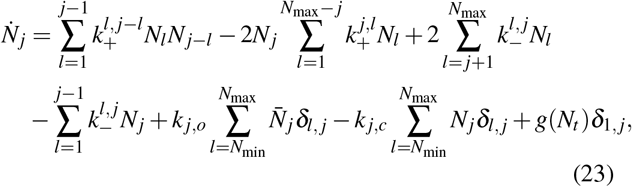

where 1 ≤ *j* ≤ *N*_max_, 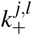 is the association rate of two oligomers of sizes *j* and *l* into an oligomer of size *j* + *l*, 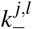 is the dissociation rate of a *j*-mer into a smaller *l*-mer and a *j*− *l* sized oligomer, *N*_*j*_ is the concentration of species *j*, 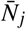 is t−he concentration of a closed ring of size *j, N*_*t*_ is the total number of adsorbed monomers, *k* _*j,c*_ and *k* _*j,o*_ are the closing and opening rates of an oligomer of size *j, N*_max_ is the maximum size of an oligomer and *N*_min_ is the minimum size of an oligomer that can form a closed ring. The first two terms are related to coagulation of two oligomers into a larger complex, whereas the two following terms quantify the fragmentation of larger structures into smaller complexes. The fourth and fifth term represent the opening and closure of species, and the last term represents the incoming flux of monomers into the surface, and hence it is added to the *j* = 1 equation exclusively. For the case of closed species, they are assumed to be formed only by closure of a previously existing open species with the same size, so their rate equations are given by

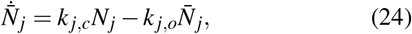

where *N*_min_ ≤ *j* ≤ *N*_max_. One can show that these equations do in fact obey mass conservation, so that

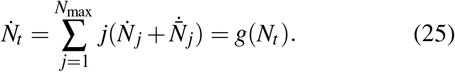

Therefore, one confirms that the term *g* is a source term for incoming monomers from the bulk. In the case of interest *g* is given by Eq. 22. A closed system may be described by Eq. 23 with *g* = 0.

Following Klein^23,25^, the rate constants 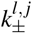 are overall rates that result from the interplay of diffusion and reaction; the reversible association of species *A* and *B* into *C* occurs in two steps,

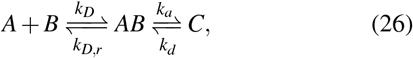

where *A* and *B* first reach reactive distance via diffusive motion with rate *k*_*D*_, thus forming a transition state *AB*. Such transition state may then react to form *C* with rate *k*_*a*_ or it may diffuse back to the reactant species with rate *k*_*D,r*_. Similarly, *C* may decay back to *AB* with rate constant *k*_*d*_. The overall rates of association and dissociation then reads,

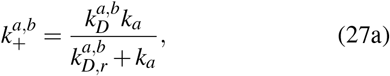

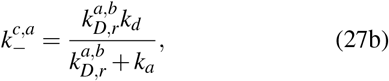

where *a, b* and *c* are the respective sizes of *A, B* and *C*. The size dependence of the constants is incorporated in the diffusive term because diffusion is dependent on the size of the species. In contrast, the reactions are mediated by identical binding sites and hence *k*_*a*_ and *k*_*d*_ are expected to be size-independent.

The limiting cases of *k*_*D,r*_≪ *k*_*a*_ and *k*_*D,r*_≫ *k*_*a*_ are known as the *diffusion limit* and *reaction limit*, respectively. For the reaction limit, the rates become

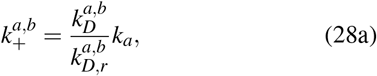

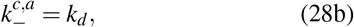

where the dissociation rate has lost its size dependence. Further simplification is achieved by assuming the forward and backward diffusion rate to be similar, 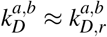, so that the overall association rate also becomes independent of size, 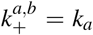. Hence, the CF equations may be written in the reaction-limited approximation as^13^

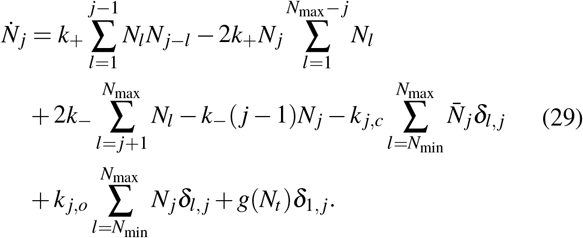

These are the macroscopic equations which will be compared to the results of the BD/GCMC routine. In particular, the main interest lies in determining the macroscopic constants as a function of the microscopic parameters *k*_±_ = *k*_±_ (*ε*_*a*_, *θ*_*c*_, …). This would allow for the quantification of the effect of the microscopic parameters on the equilibrium state of the system.

## III. RESULTS

### A. Overview

Representative snapshots of single runs of the simulation are shown in Fig. 4 (a corresponding movie is provided as supplementary material). In the following all parameters except the two most central ones, namely *ε*_*a*_ and *θ*_*c*_, are kept constant unless otherwise stated and their values are documented in the supplementary material. The snapshots qualitatively show the expected behavior of an assembly process on a surface; as the adsorption crowds the surface with free monomers (driven by the adsorption energy *V*_0_ that is held fixed at a value of 55*k*_*B*_*T* to give realistic concentrations), they are able to interact and assemble into oligomers of increasing size. As oligomers surpass a certain size, they gain the ability to close, so that for sufficiently long times one gets closed rings such as those observed in Fig. 4c and d. In the following sections, we analyze the kinetics of adsorption and assembly as well as the effect of the two central microscopic parameters *ε*_*a*_ and *θ*_*c*_. We finally compare with experimental data to estimate likely values of *ε*_*a*_ and *θ*_*c*_.

**FIG. 4:**
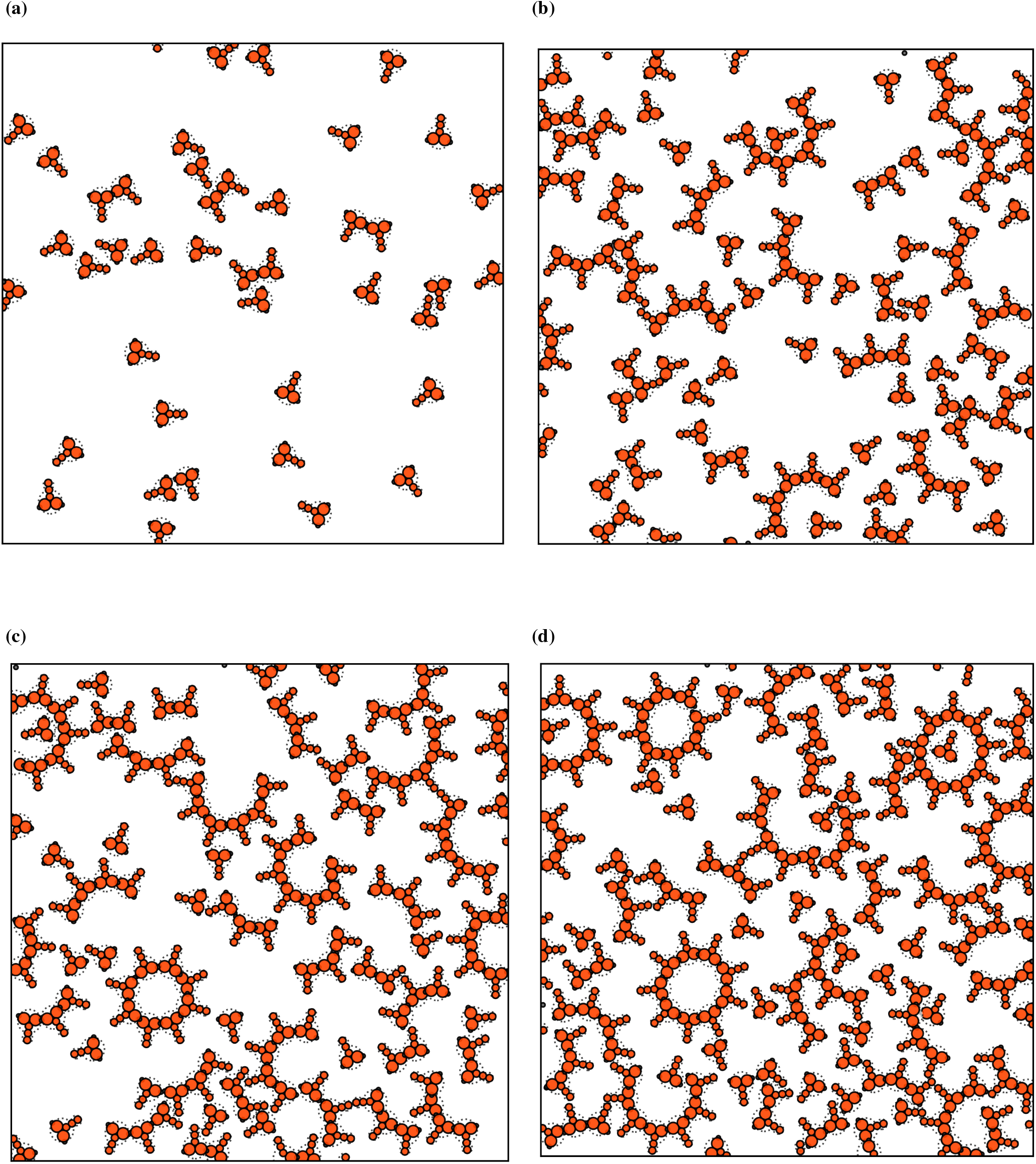
Snapshots of a simulation run at four subsequent times (a) *t*_1_, (b) *t*_2_, (c) *t*_3_ and (d) *t*_4_ for a box of size *L* = 170nm. The binding site parameters are relative binding energy *ε*_*a*_/*ε*_*r*_ = 2 and angular range *θ*_*c*_ = 0.2π. The diffusion properties are calculated from the five-bead model (shown here in red) and the binding reactions are simulated with the two-bead model (shown with dots). As the adsorption crowds the domain, oligomerization of complexes ensues. The smaller complexes are formed first and then over time give rise to larger structures. Rings exists in (c) and (d). Note that rings show deformations due to flexibility of the head-head binding, but that the spokes are straight because here we only model the short SAS-6 variant with six heptad repeats.

### B. Adsorption kinetics

Adsorption is fully specified by two main features: its steady state, which is intrinsically thermodynamic, and its kinetics. The former is described by an isotherm, which relates the final number of adsorbed particles 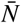 to the bulk concentration ρ, or equivalently, to the chemical potential *μ*. A simple isotherm is expected to be a monotonously increasing function that eventually saturates into a saturation limit where no additional particle may be added.

One can show that the limiting cases of Eqs. 18 in steady state coincide with these expectations, as long as the interaction potential is purely repulsive. Inserting Eq. 20 into 18, one finds that the variables *μ* and *V*_0_ act as a combined, single, effective chemical potential 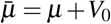. 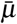 must fulfill the inequality 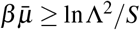 in order for at least one particle to be adsorbed on the system in steady state. For 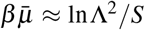, one finds that 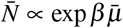. For an ideal gas, this reduces to 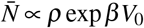. In the opposite limit of 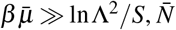 saturates, thus becoming independent of 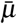. More details are given in the supplementary material. These theoretical observations are verified by simulating different values of the surface potential *V*_0_ and no assembly (*ε*_*a*_ = 0). The results are shown in Fig. 5a and imply that a reasonable adsorption energy *V*_0_ is of the order of 40*k*_*B*_*T*.

**FIG. 5:**
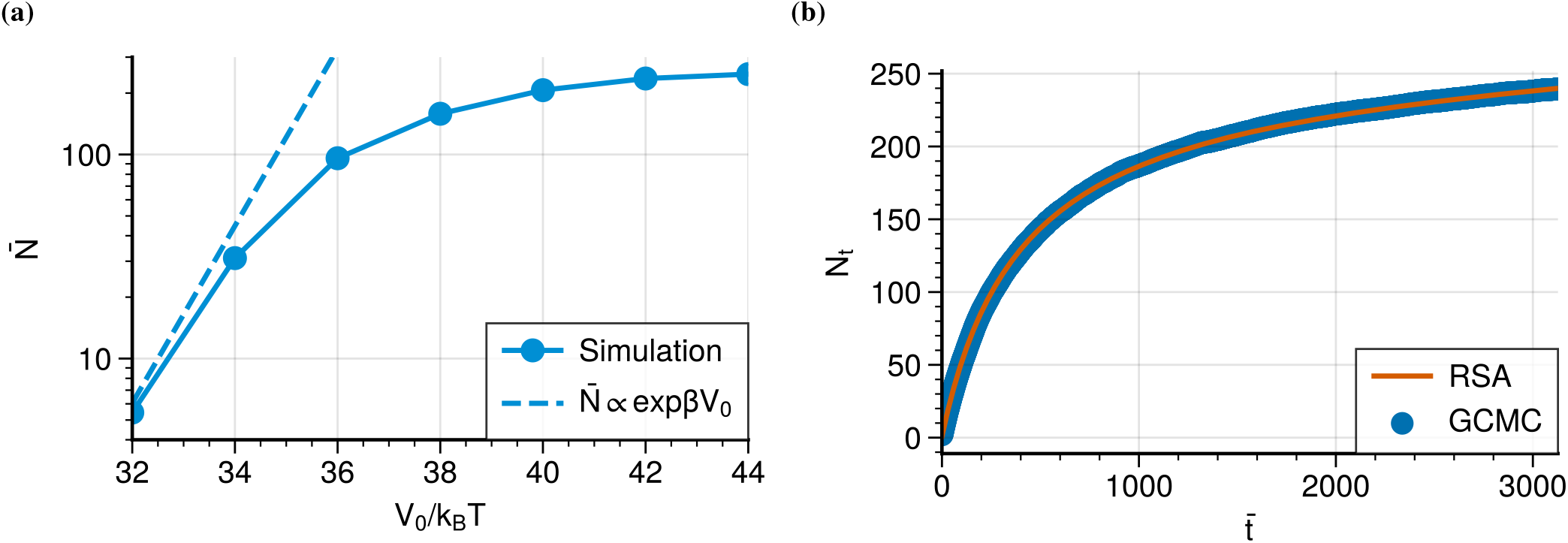
(a) Adsorption isotherm from an ideal gas reservoir with ρ = 10^23^m^−3^ onto a box with *L* = 170nm. The average final number of particles in steady state 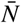 is plotted against surface potential *V*_0_. The simulated isotherm follows the expected behavior, with an increase given by 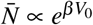 followed by saturation. (b) Comparison between GCMC simulated adsorption with *n* = 1000 and RSA kinetics with optimized parameters *k*_2_ = 0.53ns^−1^ and *k*_3_ = 1.81ns^−1^, where excellent agreement is observed. Time is rescaled as 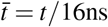.

The simulated adsorption kinetics should be well approximated by Random Sequential Adsorption kinetics, given by Eq. 22. This is because, according to Schaaf and Talbot^33^, RSA is the limiting case where the in-plane diffusion of the molecules is much slower than the adsorption rate. Therefore, the condition for RSA is that the diffusion time scale *t*_*D*_ is much larger than the adsorption time scale *t*_*A*_, so *t*_*D*_ ≫ *t*_*A*_. SAS-6 fulfills this condition; given the diffusion coefficient *D* ∼ 10^−6^cm^2^s^−1^ and the nanometric size of the protein *l* ∼ 10nm, the diffusion time scale is *t*_*D*_ ∼ *l*^2^/*D* ∼ 10^−6^s. On the other hand, the adsorption time scale is manually defined by the user through the GCMC time step in Eq. 21. In order for adsorption to be at least in the same time scale as diffusion, *n* ∼ *t*_*D*_/Δ*t* ∼ 10^5^. Hence, each GCMC step would have to be computed at least every 10^5^ time steps, thus requiring enormous computational time and resources to observe non trivial behaviour. Since in this work simulations were executed at no more than 10^6^ time steps, it follows that *n* ≪ 10^5^ and so the simulated adsorption is in the RSA limit. For the specific case of *n* = 1000 that was chosen for the multiple particle assembly simulations, Fig. 5b shows that the RSA kinetics as given by Eq. 22 with optimized parameters perfectly describe the simulated adsorption.

### C. Assembly kinetics and reaction rates

Simulations were run for 2 × 10^6^ time steps for different values of the attractive energy strength *ε*_*a*_ and cut-off angle *θ*_*c*_. The latter is chosen from the set of values {0.1π, 0.2π, 0.3π, 0.4π}. Since the assembly process will be mediated by the interplay between repulsion and attraction, the attractive energy parameter is varied as a multiple of the repulsive energy parameter *ε*_*r*_ that was kept constant at 5*k*_*B*_*T* ; the ratio *ε*_*a*_/*ε*_*r*_ is chosen from the set of possible values {1, 2, 3, 4}. For each combination of parameters 32 simulations were computed and the results were then averaged. A list of the parameters is given in the supplementary material. A first example for the results for the simulated time evolution of the concentrations is shown in Fig. 6. They are qualitatively similar to the observed kinetics in the PORT-HS-AFM experiments (compare Fig. 1f). Self-assembly proceeds in a hierarchical fashion, where small oligomers form first and are then followed by larger structures. For this particular combination of parameters, closed rings are only observed as nonamers.

**FIG. 6:**
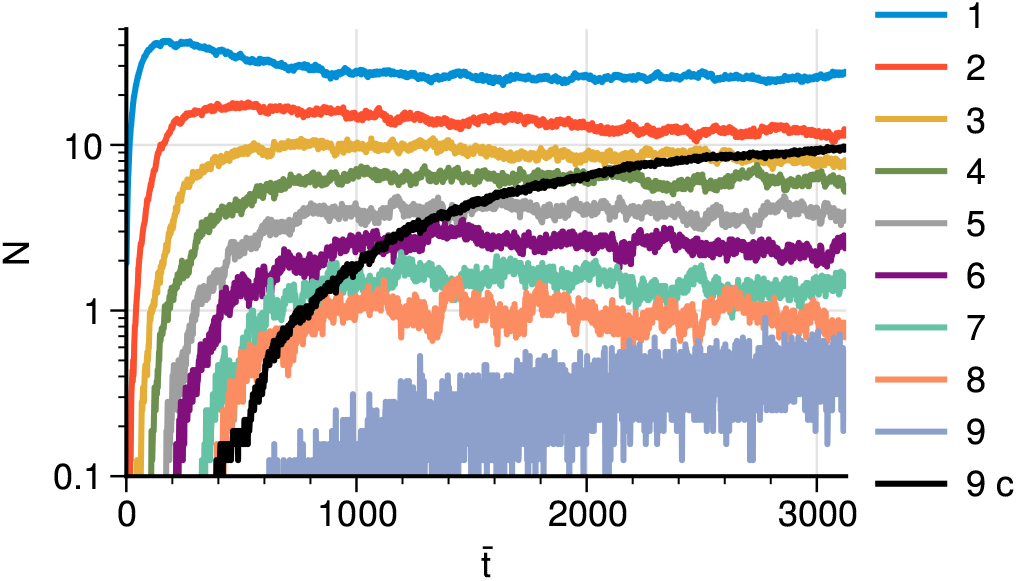
Concentrations for oligomers of different sizes as a function of rescaled time 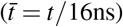 for parameters *ε*_*a*_/*ε*_*r*_ = 2 and *θ*_*c*_ = 0.1π in a box of size *L* =170nm. The time evolution is qualitatively similar to the observations in the PORT-HS-AFM experiments, where assembly occurs in a similar hierarchical fashion. The “c” label represents closed structures, which were observed to be nonamers exclusively for this combination of parameters.

One major difference between the simulation conditions and the experimental measurements is the time scale during which assembly occurs: the simulated oligomerization happens in tens of microseconds, while the experimentally observed process occurs in minutes. The explanation is that the diffusion matrix has been calculated for bulk hydrodynamics in water and did not take into account the effect of surface adhesion. The striking similarity between the simulated and observed behavior suggests that the main effect of the surface here is to dramatically slow down the timescale, but does not change the relative importance of the different sub-processes. We note that in general it is challenging to improve on the hydrodynamic model, because the details of the interaction with the surface are not known.

We next turn to the reaction rates and analyze the concentration profiles to the CF equations in the reaction limited approximation, as given by Eq. 29. The parameters *k*_±_ and *k*_*o,c*_ are optimized by minimizing the average relative square error between the simulated concentrations and the numerical solution of Eq. 29. Such minimization was done on a case by case basis because each combination of parameters (*ε*_*a*_, *θ*_*c*_) resulted in unique concentration profiles with different minimum closing size *N*_min_ and maximum oligomer size *N*_max_; fitting by brute force all the curves to a single combination of *N*_min_ and *N*_max_ would be non-physical and lead to poor estimation of the rate parameters. A full recollection of the fitting results is found in supplementary material.

In Fig. 7 we compare the results of our computer simultions with the numerical solution of the CF-equations. For an energy ratio of *ε*_*a*_/*ε*_*r*_ = 1 (Fig. 7a), the attractive potential is too weak to stabilize large oligomers. As a consequence, the majority of monomers remain unbound at the end of the simulation, and only oligomers no larger than tetramers are able to assemble in appreciable quantites. This is reflected in the CF-equations by setting *N*_max_ = 4 and neglecting all terms related to closed structures. Moreover, an increase in the angle parameter (Fig. 7b) favors association of oligomers by amplifying the angular range of the attractive potential, so all the concentration curves are shifted upwards, except for the monomers. This effect is nevertheless insignificant, as most monomers remain free even for the largest cutoff *θ*_*c*_ = 0.4π. The angular cutoff therefore plays a secondary role at this energy range.

**FIG. 7:**
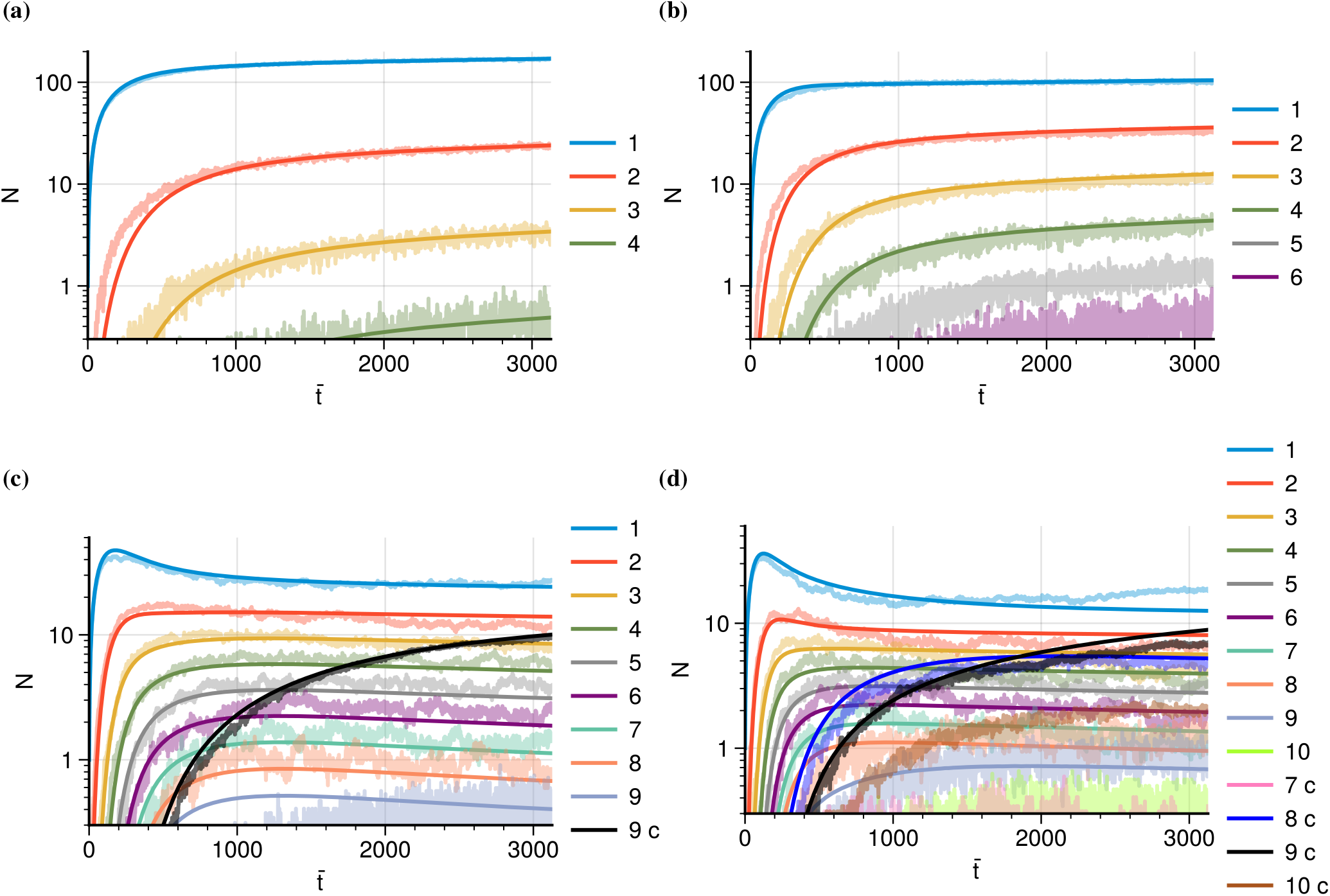
Concentration of oligomers of different sizes in a box of size *L* = 170nm as a function of rescaled time 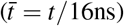 for (a) *θ*_*c*_ = 0.1π and *ε*_*a*_/*ε*_*r*_ = 1, (b) *θ*_*c*_ = 0.4π and *ε*_*a*_/*ε*_*r*_ = 1, (c) *θ*_*c*_ = 0.1π and *ε*_*a*_/*ε*_*r*_ = 2 and (d) *θ*_*c*_ = 0.3π and *ε*_*a*_/*ε*_*r*_ = 2. The results of the simulations, the light colored curves, are compared to the numerical solution of the CF-equations, the darker overlaid lines, with optimized parameters *k*_±_. The label “c” indicates a closed species. Decamers in Fig. (d), as well as pentamers and hexamers in Fig. (b), are shown but not considered in the CF comparison. Species with concentrations below N = 0.3 are considered negligible and thus not shown.

In contrast, for *ε*_*a*_/*ε*_*r*_ = 2 (Fig. 7c and d) full assembly is observed, so *N*_max_ = 9. Higher order structures such as decamers, which assemble in limited quantities, are therefore neglected in the CF-equations. As seen in Figs. 7c (*θ*_*c*_ = 0.1π) versus 7d (*θ*_*c*_ = 0.3π), the system is much more sensitive to the angular cutoff *θ*_*c*_ for *ε*_*a*_/*ε*_*r*_ = 2. Not only does this parameter affect the interplay between dissociation and association, it also mediates the survival or extinction of entire populations of closed oligomers. For a restrictive choice of *θ*_*c*_ = 0.1π the formation of malformed structures is suppressed at all times. It follows that the minimum closed size must be set *N*_min_ = 9 in order for the CF equations to properly describe the dynamics. A more permissive angular cutoff such as *θ*_*c*_ = 0.3π allows greater angular deviations and so malformed rings such as octamers become possible. Hence, the minimum closed ring is *N*_min_ = 8, and this is also the case for *θ*_*c*_ = 0.2π and *θ*_*c*_ = 0.4π.

In Fig. 8 we compare simulations and CF-equations for higher values of the energy ratio, namely *ε*_*a*_/*ε*_*r*_ = 3 and 4. These two energies show similar dynamics, so the parameters *N*_max_ and *N*_min_ are solely determined by the angle cutoff *θ*_*c*_. For the lowest value of *θ*_*c*_, only closed nonamers were observed, so *N*_max_ = 9 and *N*_min_ = 9. However, no open nonamers are observed in significant quantities; the closure of the nonamer has become *instantaneous*. Any oligomer with nine proteins will immediately form a ring, so the transient open species is short lived. This is a limiting case of the CF equations, where *k*_9,*c*_/*k*_9,*o*_ ≫ 1. Instantaneous closure is already visible in Fig. 7c for an energy ratio of 2. The CF equations overestimate the concentration of the open nonamers, suggesting that the closure rate is faster than estimated. Microscopically, this occurs because the combination of a strong attractive potential and a narrow angular range result in a stiff bond which hardly deforms from its equilibrium angle ϕ_*eq*_ = 40°. The interior angles of any nonamer will not deviate significantly from those of a perfect nonagon, so any nine-sized complex will immediately close. This fact may be captured in the CF equations by eliminating the terms of closure and opening of nonamers, and replacing the open concentration with its closed counterpart 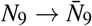 in the remaining equations.

**FIG. 8:**
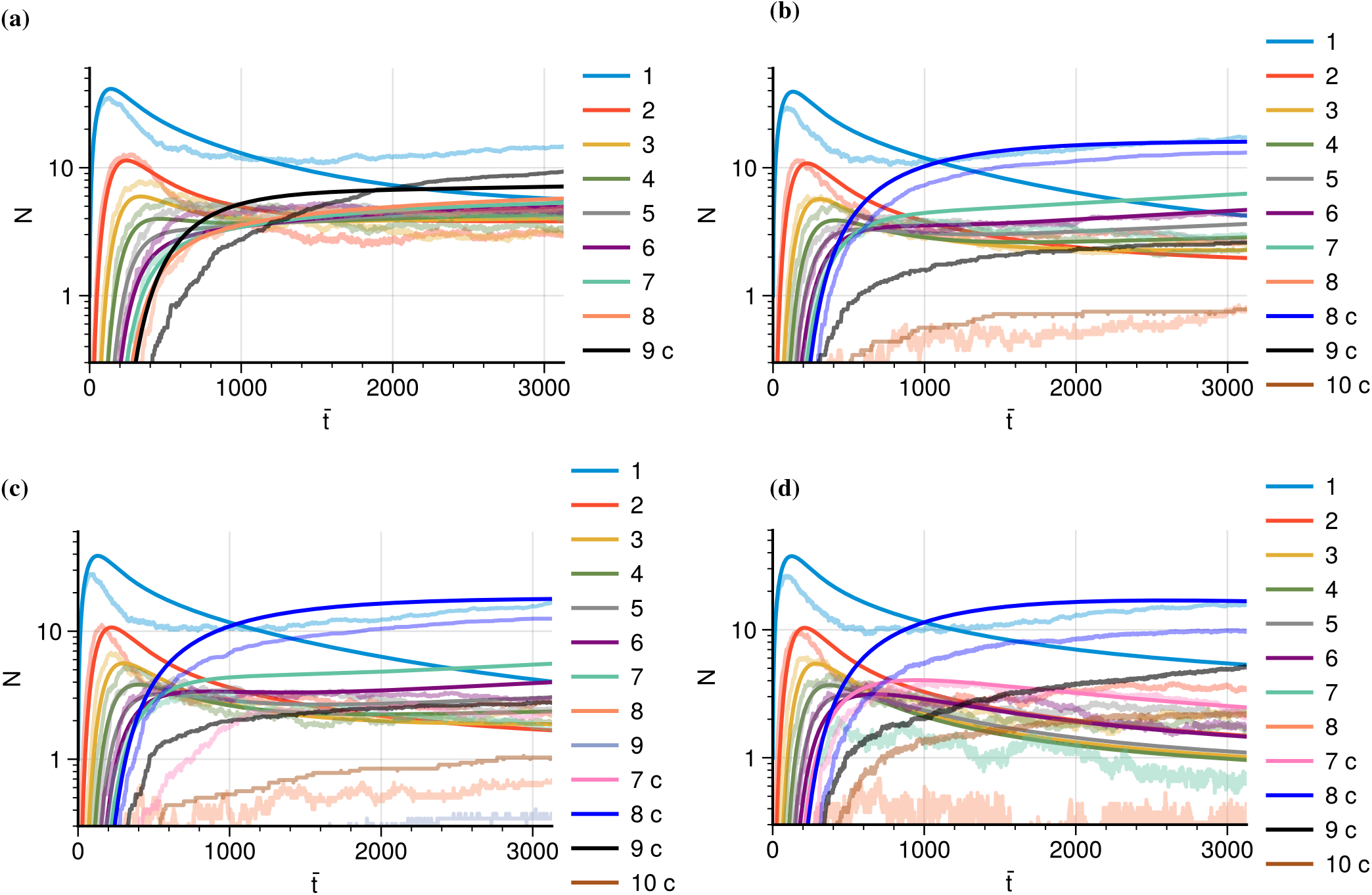
Concentration of oligomers of different sizes in a box of size *L* = 170nm as a function of rescaled time 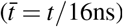 for (a) *θ*_*c*_ = 0.1π and *ε*_*a*_/*ε*_*r*_ = 3, (b) *θ*_*c*_ = 0.2π and *ε*_*a*_/*ε*_*r*_ = 4, (c) *θ*_*c*_ = 0.3π and *ε*_*a*_/*ε*_*r*_ = 4, and (d) *θ*_*c*_ = 0.4π and *ε*_*a*_/*ε*_*r*_ = 3. The results of the algorithm, the light colored curves, are compared to the numerical solution to the CF equations, the darker overlaid lines, with optimized parameters *k*_±_. Light colored curves without a darker counterpart were neglected in the CF comparison. For *θ*_*c*_ = 0.2π and 0.3π only octamers are included as closing species in the CF comparison, whereas for *θ*_*c*_ = 0.4π closed heptamers are also included. Species with concentrations below N = 0.3 are considered negligible and thus not shown.

For larger cutoff values, the formation of nine-sized rings is severely suppressed and malformed structures become dominant. The octamer has become the closed species with the largest relative population. In the CF equations, the predominant species is implicitly determined by *N*_max_; in a system where association is favourable, oligomers will keep coagulating into larger structures until they stop at *N*_max_. It follows that *N*_max_ = 8 for *θ*_*c*_ ≥ 0.2π, which was the choice with the better fit. Furthermore, no significant quantities of smaller sized complexes are observed for *θ*_*c*_ = 0.2π and *θ*_*c*_ = 0.3π, so *N*_min_ = 8. However, closed heptamers become important for *θ*_*c*_ = 0.4π, so *N*_min_ = 7. For all cases the closure of rings is approximated as instantaneous.

The main observation in Fig. 8 is that, compared to lower energy ratios, the CF equations are no longer able to describe the time evolution of the system for *ε*_*a*_/*ε*_*r*_≥ 3. In all cases the coagulation fragmentation equations predict a faster than observed depletion of free monomers and small sized oligomers.

The discrepancies are particularly evident for increasing *θ*_*c*_; in this case, the population of closed octamers is overestimated at all times, and larger structures such as the closed nonamer and decamer could not be accounted for in the best fit. This suggests that the reaction limited approximation is no longer valid in the range of high attractive energy parameter and large cutoff angle. The wide ranged potential is strong enough to rapidly bond two monomers as soon as they are within reactive distance. Therefore, the limiting step of the association events is the diffusive motion to actually reach this reactive threshold, and so the system becomes diffusion limited.

### D. Efficiency and selectivity

In order to qualitatively assess the effect of these microscopic parameters on the state of the system at the end of the simulation, two metrics are defined: the final fraction of monomers that are bound to a closed species (*efficiency*) and the population of the biologically desired closed nonamers relative to the population of all closed complexes (*selectivity*). The former is given by the bound fraction ξ and the latter is the selectivity of the nonamer *S*_9_, which are mathematically defined as

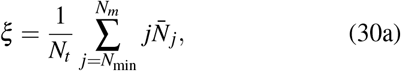

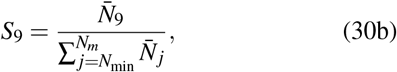

where *N*_*t*_ is the total number of monomers on the surface, 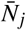 is the number of rings with *j* proteins, *N*_min_ is the minimum closing size and *N*_*m*_ is the maximum observed size of the closed oligomers. The first metric quantifies how efficient is the formation of closed structures compared to intermediate open structures, whereas the second parameter measures how selective the system is for the formation of closed nonamers in comparison to rings of different sizes.

Our results for efficiency and selectivity are shown in Fig. 9a and b, respectively. Increasing *θ*_*c*_ and *ε*_*a*_ leads to a more numerous population of closed structures and a higher fraction of monomers bound to a ring (Fig. 9a). This is understandable from a physical perspective: higher values of *ε*_*a*_ lead to a deeper potential well, so dissociation of complexes becomes unlikely and the formation of larger, closed structure is facilitated. In addition, larger values of *θ*_*c*_ increase the *E*_*a*_ number of available bonding candidates, given that monomers within a larger span of relative orientations will experience an attractive potential, and so association is expedited.

**FIG. 9:**
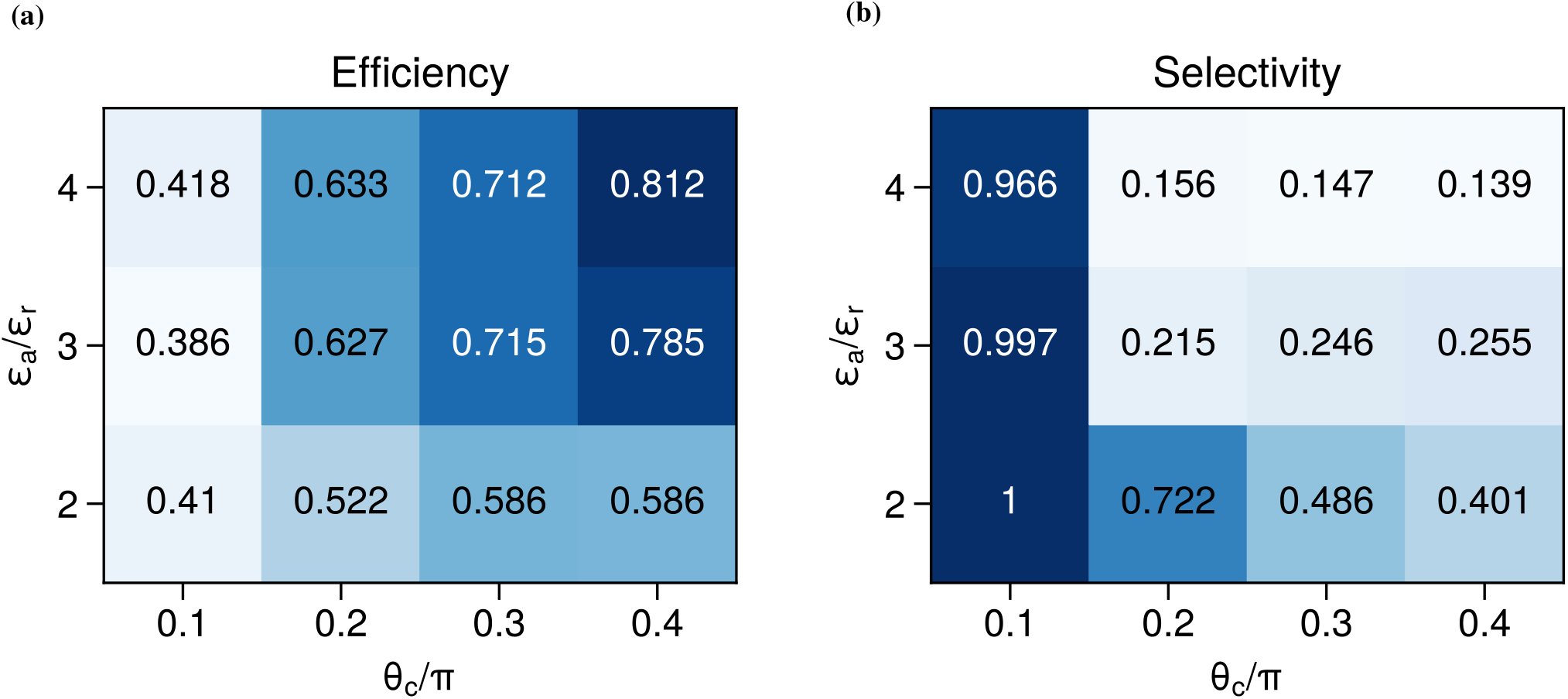
(a) Efficiency as measured by the fraction of monomers ξ which are bound in a closed complex. Higher values of *ε*_*a*_/*ε*_*r*_ and *θ*_*c*_ result in a large fraction of monomers bound to a ring, at the expense of the formation of malformed structures. (b) Selectivity as measured by the relative population *S*_9_ of closed complexes that exhibit nine-fold symmetry. Larger values of the angular cutoff *θ*_*c*_ lead to the formation of malformed structures and thus decrease *S*_9_. For energy ratio *ε*_*a*_/*ε*_*r*_ = 1, no closed rings were observed.

However, the larger bound fraction comes at the expense of a higher relative population of malformed complexes (Fig. 9b). For larger values of the attractive strength and the angular cutoff, nonamers only constitute 10%-25% of the total number of closed complexes and must be neglected completely in order to obtain a proper fit of the CF equations. In contrast, for the lowest value of *θ*_*c*_ = 0.1π nonamers compose more than 95% of all closed complexes. Such a sharp difference in nonamer assembly has two underlying reasons. The first one has to do with the angular cutoff *θ*_*c*_. An increase of this parameter results in larger deviations from perfect alignment, corresponding to a nonagon, so that malformed structure become possible. This is why the most restrictive value of *θ*_*c*_ = 0.1π results in the highest observed selectivities. The second, and somewhat more subtle reason, has to do with the attractive energy strength. A malformed structure may, in principle, form a desired nonamer by breaking bonds and bonding with additional oligomers, if it has less than nine proteins, or fragmenting into smaller complexes if it has more. These processes should be thermodynamically desired because the nine protein ring is by construction the minimum energy configuration. However, the step of bond breaking is key for such process to happen; bond formation must be *reversible*. If the potential well is too high, as it is the case of energy ratios *ε*_*a*_/*ε*_*r*_ = 3 and 4, bond breaking is unlikely and any malformed structure will remain intact, even if there exists a lower energy configuration. This phenomenon is known in the literature as *kinetic trapping* and it explains why the energy ratio *ε*_*a*_/*ε*_*r*_ = 2 is less prone to the formation of malformed structures in comparison to its higher counterparts.

An energy ratio of 2 is sufficiently low for bond reversibility and thus the nonamer formation is favoured. Energy ratios of 3 and 4, by contrast, are in a regime were kinetic trapping is dominant, and thus they exhibit much lower selectivites to the nonamer formation.

### E. Comparison with experiments

The estimated parameters *k*_±_ allow for the quantification of the dissociation constant *K*_*D*_, defined as,

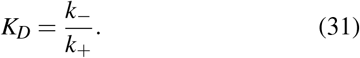

The dissociation constant is an experimentally measurable quantity of the thermodynamic balance between dissociation and association. The results from our simulations are shown in Fig. 10a. We see that ln *K*_*D*_ exhibits a rough linear decay with energy ratio *ε*_*a*_/*ε*_*r*_ in the range *ε*_*a*_/*ε*_*r*_≤ 3. Furthermore, the slope seems to be unaffected by the angular cutoff *θ*_*c*_, so that it only influences the intercept with the vertical axis. These are the expected features of a thermodynamic equilibrium constant that follows an Arrhenius-type equation

**FIG. 10:**
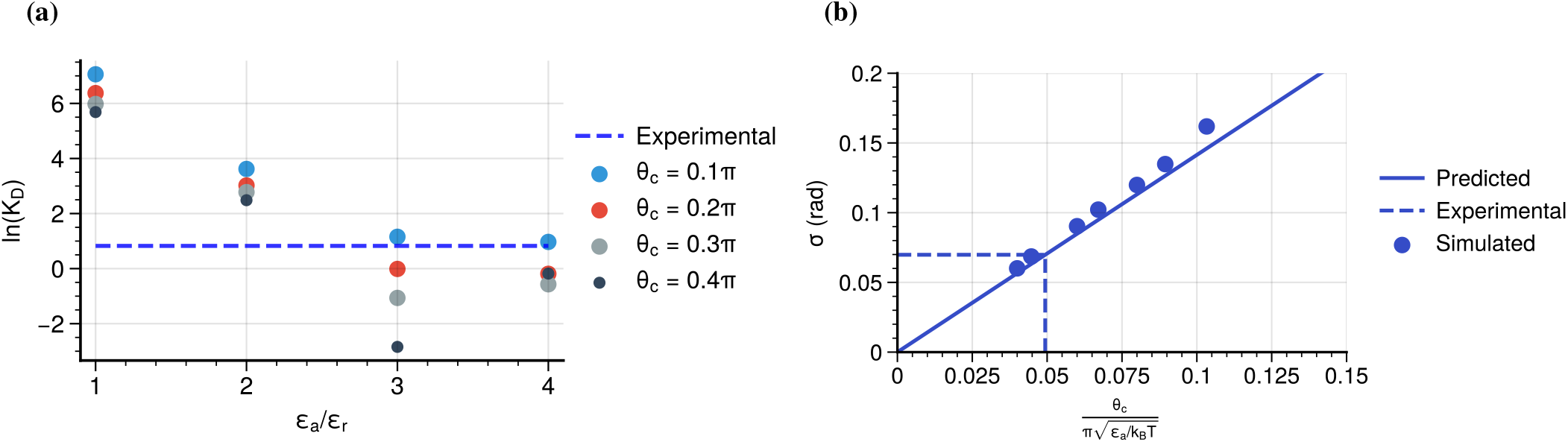
(a) Dissociation constant *K*_*D*_ as a function of energy ratio *ε*_*a*_/*ε*_*r*_ and cutoff angle *θ*_*c*_. For *ε*_*a*_/*ε*_*r*_≤ 3, the dissociation constant follows roughly an exponential decay with a preexponential factor determined by the angular parameter. The dashed blue line indicates the experimental *K*_*D*_ on mica as reported by Banterle^13^. (b) Bond angle standard deviation of a single bond as a function of *θ*_*c*_ and *ε*_*a*_. To lowest order, the angular fluctuations of a bonded pair are proportional to the ratio 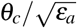, something which is confimed by the simulations. The dashed black line labels the experimental standard deviation of 4°reported by Banterle^13^ for a tetramer, which corresponds to 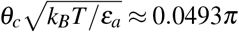.

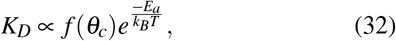

where *E*_*a*_ is the free activation energy of the reaction and *f* (*θ*_*c*_) is the steric factor accounting for bonding anisotropy. Beyond *ε*_*a*_/*ε*_*r*_ = 3 the dissociation constant saturates, confirming that the system is now diffusion limited; any further increase in the attractive energy parameter will no longer decrease the equilibrium constant as the slowest step is related to diffusive motion and not to the chemical assembly. The estimation of *K*_*D*_ in the high energy range would require going beyond the reaction limited approximation of the CF equations and consider rates of diffusion, which would inevitably lead to size dependent rate constants 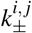.

Experimentally, Banterle obtained *K*_D_ = 79*μ*m^-2^ for oligomerization on a surface^13^, corresponding to a value of 2.28 for our box size of 170nm (dashed line in Fig. 10a). One sees that our simulations give similar values between *ε*_*a*_/*ε*_*r*_ = 2 and 3 (depending on *θ*_*c*_). This is also the regime in which assembly already gives many closed complexes (in contrast to *ε*_*a*_/*ε*_*r*_ = 1, where one mainly gets small oligomers). At the same time, this is also the regime in which ring opening and closure still occur, and in which the CF-equations still work, as also observed by Banterle. This suggests that this is the range which corresponds to the experiments.

This conclusion also agrees with the selectivities experimentally observed by Hilbert^17^, where 43 % of the closed structures are nonamers. In Fig. 9b, one observes a sharp decrease between 99% and 20% between 0.1π ≤ *θ*_*c*_ < 0.2π for *ε*_*a*_/*ε*_*r*_ = 3, so 43% should lie somewhere in between. From these considerations we conclude that realistic parameter values for the PORT-HS-AFM experiments should lie in the range 0.1π ≤ *θ*_*c*_ < 0.2π and 2 < *ε*_*a*_/*ε*_*r*_ ≤ 3.

To better constrain the experimentally relevant parameter values, we finally consider the fluctuations of the bond angles. One can approximate the bond angle dynamics by a harmonic oscillator by expanding Eq. 9 to lowest order around the minimum. In the presence of Gaussian noise, this becomes an Ornstein Uhlenbeck stochastic process, whose steady state probability distribution is a Gaussian with a variance σ^2^ that is inversely proportional to the potential strength (the repulsive potential strength *ε*_*r*_ is not relevant in this context):

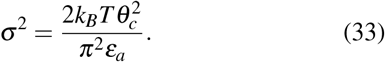

We simulated a single bond for different combinations of *ε*_*a*_ and *θ*_*c*_ and confirmed that the simulations do indeed follow Eq. 33 (Fig. 10b). The bounds for *ε*_*a*_ and *θ*_*c*_ suggested above lead to a theoretical interval of 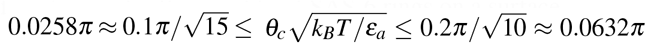. This matches the experimental value reported by Banterle^13^ of a standard deviation of 4°for a tetramer (dashed line in Fig. 10b). T he experimental result corresponds to 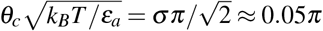, thus validating the fact that the microscopic parameters are within the proposed range. A reasonable combination that reproduces the observed variance would be *ε*_*a*_/*ε*_*r*_ = 2.2 and 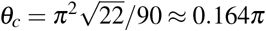.

## IV. DISCUSSION

Here we have developed and implemented a GCMC/BD algorithm for adsorption and 2D ring self-assembly of patchy particles representing SAS-6 dimers. Our BD-approach allows us to simulate the whole assembly process and at the same time to also study the effect of microscopic parameters like diffusion and rate constants. In contrast to earlier BD-simulations for SAS-6 ring assembly in solution^25^, which modeled the patches by their geometrical effect and not by potentials as done here, our approach allows us to explore the formation of malformed closed complexes. The importance of this feature has become evident by the PORT-HS-AFM experiments that revealed that besides the expected nine-rings, also seven, eight and ten rings can form, revealing a high degree of conformational plasticity in the SAS-6 system. By varying the microscopic binding parameters *ε*_*a*_ and *θ*_*c*_, we can predict the frequency of these different structures. Our computer simulations also allowed us to study transitions between the diffusion and reaction limits, the role of bond anisotropy and the effect of kinetic trapping. Furthermore, the simulations provide a hint towards surface induced energy reduction of association reaction, presumably by restricting the ability of the oligomers to bend. These observations might help elucidate the assembly of the centriole in a cellular context, as well as the role of the cartwheel structure in the biological function of the centriole and the reason for its evolutionary conservation.

The main findings of our simulations may be summarized as follows: in general the model is able to qualitatively reproduce the time evolution of oligomer concentration as observed in PORT-HS-AFM experiments. For low energy ratios *ε*_*a*_/*ε*_*r*_ ≤ 2, the system is well described by the CF equations in the reaction limit approximation. In contrast, higher energy ratios *ε*_*a*_/*ε*_*r*_ ≥ 3 no longer agree with this regime, as the strong attractive potential causes the system to become diffusion-limited. Regarding the equilibrium state, we find that the energies and angular cutoffs mediate the interplay between bound fraction and selectivity, which behave mutually exclusively (Fig. 11). Lower energy ratios and angular parameters result in high selectivity of the nonamer due to highly anisotropic bonding and reversibility, at the expense of lower bound fractions. Conversely, higher energy ratios and angular parameters preclude dissociation and widen the range of the attractive potential, thus attaining large bound monomer fraction at the cost of large relative population of malformed structures. Our quantitative comparison between simulations and experiments suggest that the SAS-6 system is positioned at the sweet spot where it can achieve both high efficiency (through relative weak binding energies) and high selectivity for the nine-fold symmetry (through a small angular range).

**FIG. 11:**
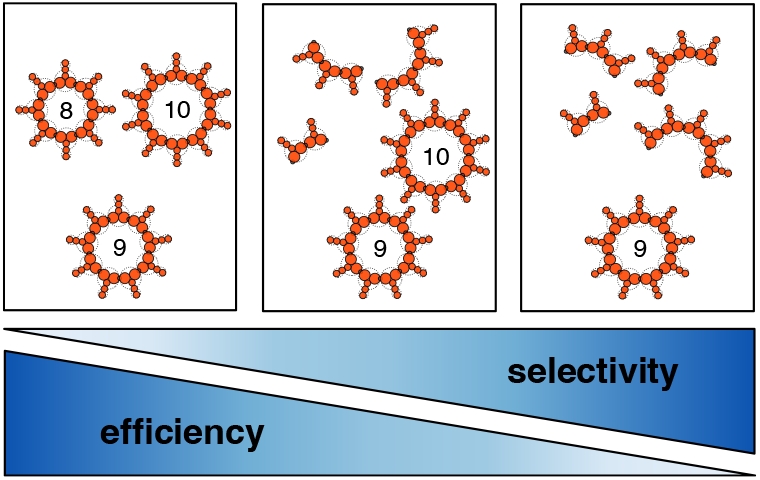
SAS-6 oligomerization is a balancing act between bonding efficiency and selectivity; either efficiency of closed complex formation is favoured at the cost of having malformed structures, or the selectivity of nonamer formation is favoured at the cost of efficiency.

A major limitation of our model is that it does not predict the correct time scale for assembly as observed in the PORT-HS-AFM experiments. In the simulations, this timescale is microseconds, as determined by the viscosity of water and bulk hydrodynamics. In the experiments, it is minutes, as determined by the interactions of the proteins with the mica surface. This dramatic slowdown can be explained by several factors, including electrostatic interactions with patches on the mica, topographical barriers and the high effective viscosity of water in small spaces (as known from lubrication analysis). In the future, the hydrodynamic model could be updated by a more realistic one, e.g. by adapting the Frenkel-Kontorova-Tomlinson model for nanoscale friction^34^. At the current stage, however, we do not have sufficient information to define a suitable model for this aspect. Yet we note that apart from the time scale, our simulations give very similar results as the experiments, suggesting that the relative importance of the different sub-processes is not changed by the interactions with the surface.

An interesting direction for further research will be to analyze the influence of the spoke length on the assembly process. The encounter of two proteins is possible by either diffusive motion of proteins on the surface or by adsorption of a protein near an already existing protein. Both of these effects are hindered by a larger protein; friction increases with increasing size, as illustrated by Stokes equation *D* ∼ 1/*r*. Hence, the diffusive motion should be restricted for larger proteins and so the encounter of two proteins happens at a slower rate. Furthermore, rotational diffusion would also suffer a reduction, which would further decrease the association rate given the anisotropy of the bonding. Additionally, a larger protein occupies a larger area on the surface, so the overall rate of adsorption also decreases because there is an augmented fraction of exclusion area per protein. However, from the cellular point of view, it is important that the cartwheel is sufficiently large to seed the microtubule part of the centriole, thus a large spoke length is required. A solution to this problem might be some degree of flexibility in the coiled-coil that could have evolved a specific disruption of its heptad repeat. Suchflexibility could be complemented by strain in the ring; together these elements might be important to build up the large-scale structure of the centriole^35^.

Our simulations demonstrate that in the case of SAS-6 self-assembly, efficiency of the assembly process and nonamer selectivity cannot be fully achieved at the same time; tuning the parameters to increase either inevitably leads to the decrease of the other. This is consistent with findings of *in vitro* experiments of SAS-6 on mica^17^, where free monomers were observed in negligible quantities and 43% of the closed structures were nonamers. However, the picture seems to be different in a cellular context, where high bound fractions and high selectivity to the nonamer were observed^35^. Such discrepancy suggests that in cells additional mechanisms exists to correct malformed structures. A proposed mechanism is the binding of additional proteins to SAS-6 dimers. In particular, the proteins Plk4 and STIL have been found to cooperate with SAS-6 at the onset of centriole assembly by focusing the SAS-6 oligomers on the resident centriole^35^. It is hypothesized that the interactions between Plk4, STIL and SAS-6 is the reason behind finely tuned ring formation. An important role is also played by Bld10p/Cep135, which links the cartwheel to the microtubule triplets^36^. In principle, the BD-framework established here can also be used to achieve a better understanding of the function of these proteins in the assembly process; by including them in the simulation, one may learn about their potential role in correcting malformed structures, stacking the SAS-6 oligomers into the cartwheel and connecting them to the microtubules. Consequently, the hereby developed routine may also shed light on how these proteins cooperate in cells in the construction of one of the key eukaryotic organelles.

## SUPPLEMENTARY MATERIAL

The supplementary material contains four supplementary figures, three supplementary tables and one movie corresponding to the simulation snapshots from Fig. 4.

## ACKNOWLEDGMENTS

We acknowledge funding through the Max Planck School Matter to Life, supported by the German Federal Ministry of Education and Research (BMBF) in collaboration with the Max Planck Society; and by the Heidelberg-Karlsruhe Cluster of Excellence 3DMM2O, supported by the Deutsche Forschungsgemeinschaft (DFG, German Research Foundation) under the Excellence Strategy (EXC 2082/1-390761711). USS is a member of the Interdisciplinary Center for Scientific Computing (IWR) at Heidelberg. We thank Falko Ziebert, Svenja de Buhr, Frauke Gräter and Camilo Aponte for stimulating discussions.

## CONFLICT OF INTEREST

The authors have no conflicts to disclose.

## DATA AVAILABILITY STATEMENT

The source code for our simulations is available on GitHub at https://github.com/sgomezmelo/Brownian-Dynamics.

## Notes

### Competing Interest Statement

The authors have declared no competing interest.

