## Supplementary figures and tables for "Grand canonical Brownian dynamics simulations of adsorption and self-assembly of SAS-6 rings on a surface"

#### A Interaction Potentials

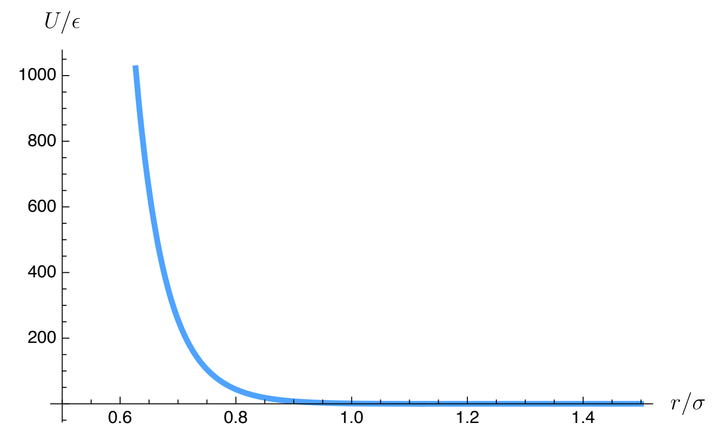


Figure S1: Repulsive potential as a function of distances between the centers of two spheres.

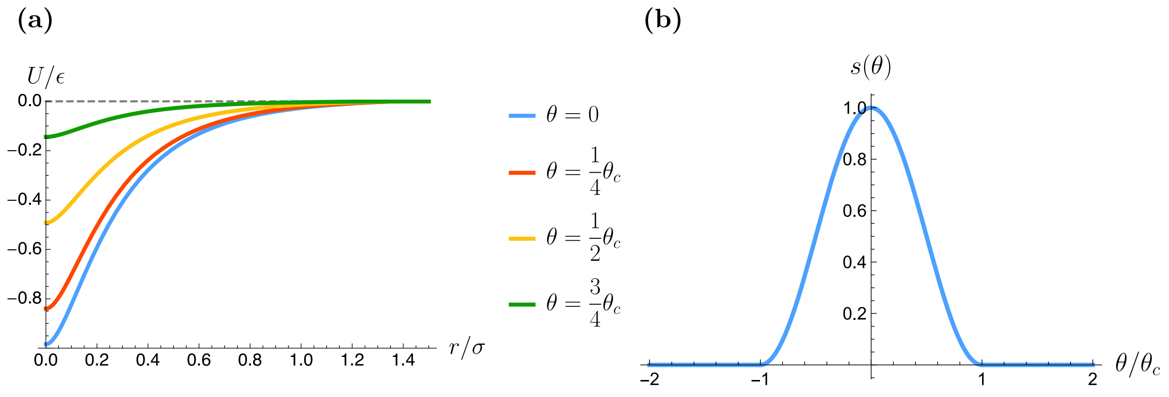


Figure S2: Dependence of the binding potential on (a) the distance between binding sites, for different bond orientiations and (b) the bond angle.

#### B Adsorption Kinetics

To observe that the Metropolis Hastings algorithm does recover the form of an isotherm in steady state, we start by demanding that in such state the rates of adsorption and desorption are equal,

$$\frac{\bar{N}}{S\Lambda^{-2}} e^{\beta\Delta U_I(\bar{N})-\beta(\mu+V_0)} = 1, \tag{1}$$

where $\bar{N}$ is the final number of particles, averaged over spatial configurations. We consider a strictly repulsive potential $\Delta U_I \geq 0$. In order to obtain at least one particle in steady state $\bar{N} \geq 1$, one must have

$$\mu + V_0 \geq \mu + V_0 - \Delta U_I \geq \frac{1}{\beta} \ln \frac{\Lambda^2}{S}, \tag{2}$$

Now consider the two limiting cases. For $\beta\bar{\mu} \to \ln \frac{\Lambda^2}{S}^+$, $\bar{N} \to 1$ so $\Delta U_I \to 0$, from which it follows that

$$\bar{N} = \Lambda^{-2} S e^{\beta\bar{\mu}} \Rightarrow \bar{N} \propto e^{\beta\bar{\mu}}. \tag{3}$$

In particular, for an ideal gas, one finds

$$\bar{N} = \rho\Lambda S e^{\beta V_0} \Rightarrow \bar{N} \propto \rho e^{\beta V_0}. \tag{4}$$

For the converse limit, one must first prove that the final number of particles $\bar{N}$ is an increasing function of the effective chemical potential $\bar{\mu} = \mu + V_0$. To do so, one simply needs to check that the derivative of $\bar{N}$ is strictly positive. This is the case,

$$\frac{d\bar{N}}{d\bar{\mu}} = \frac{\Lambda^{-2} S\beta e^{\beta(\bar{\mu}-\Delta U_I)}}{1 + \beta\bar{N}\frac{dU}{d\bar{N}}} \geq 0, \tag{5}$$

because the potential is repulsive, which implies,

$$\frac{dU_I}{d\bar{N}} \geq 0. \tag{6}$$

In other words, work is required to add any additional particle. Consequently, in the limit $\beta\bar{\mu} \gg \ln \frac{\Lambda^2}{S}$, one has a high number of particles at a distance $\sim 1/\sqrt{\bar{N}}$. It follows that the interaction term $\Delta U_I$ becomes dominant, so

$$\bar{N} = S\Lambda^{-2} e^{\beta\Delta U_I(\bar{N})}, \tag{7}$$

where it is now clear that the final particle number plateaued at a value no longer dependent on $\bar{\mu}$. This completes the argument that the Metropolis Hastings algorithm should recover the form of the isotherm.

### C Assembly Kinetics

#### C.1 Simulation parameters

The following parameters were kept constant throughout all assembly kinetics simulations.

| Parameter | Value |
|---|---|
| Body-body repulsive cutoff $c_{bb}$ (nm) | 8.2 |
| Tail-tail repulsive cutoff $c_{tt}$ (nm) | 3.5 |
| Body-tail repulsive cutoff $c_{tb}$ (nm) | 5.9 |
| Repulsive energy strength $\epsilon_r/k_BT$ | 5.0 |
| Surface energy strength $V_0/k_BT$ | 55.0 |
| Domain box length $L$ (nm) | 170.0 |
| Initial number of particles $N_0$ | 1 |
| Reservoir density $\rho$ ($\mathrm{m}^{-3}$) | $10^{23}$ |
| Temperature $T$ (°C) | 20.0 |
| Maximum bonded distance $r_b$ (nm) | 2.5 |
| Molecular weight (Da) | 51628.52 |
| Time step $\Delta t$ (ps) | 25 |
| GCMC time step $\Delta t_{\mathrm{GCMC}}$ | 1000 $\Delta t$ |

Table S1: Constant parameters in the multiple particle simulations.

#### C.2 Simulation Results

The multiple assembly simulations yielded the following concentration profile results for the parameter scan of $\epsilon_a/\epsilon_r \in \{1, 2, 3, 4\}$ and $\theta_c \in \{0.1\pi, 0.2\pi, 0.3\pi, 0.4\pi\}$, shown in figures S3 and S4. Overlaid lines are the numerical solution of the coagulation fragmentation (CF) equations with optimized parameters $k_\pm$ that minimize the total squared error, which are reported in table S2.

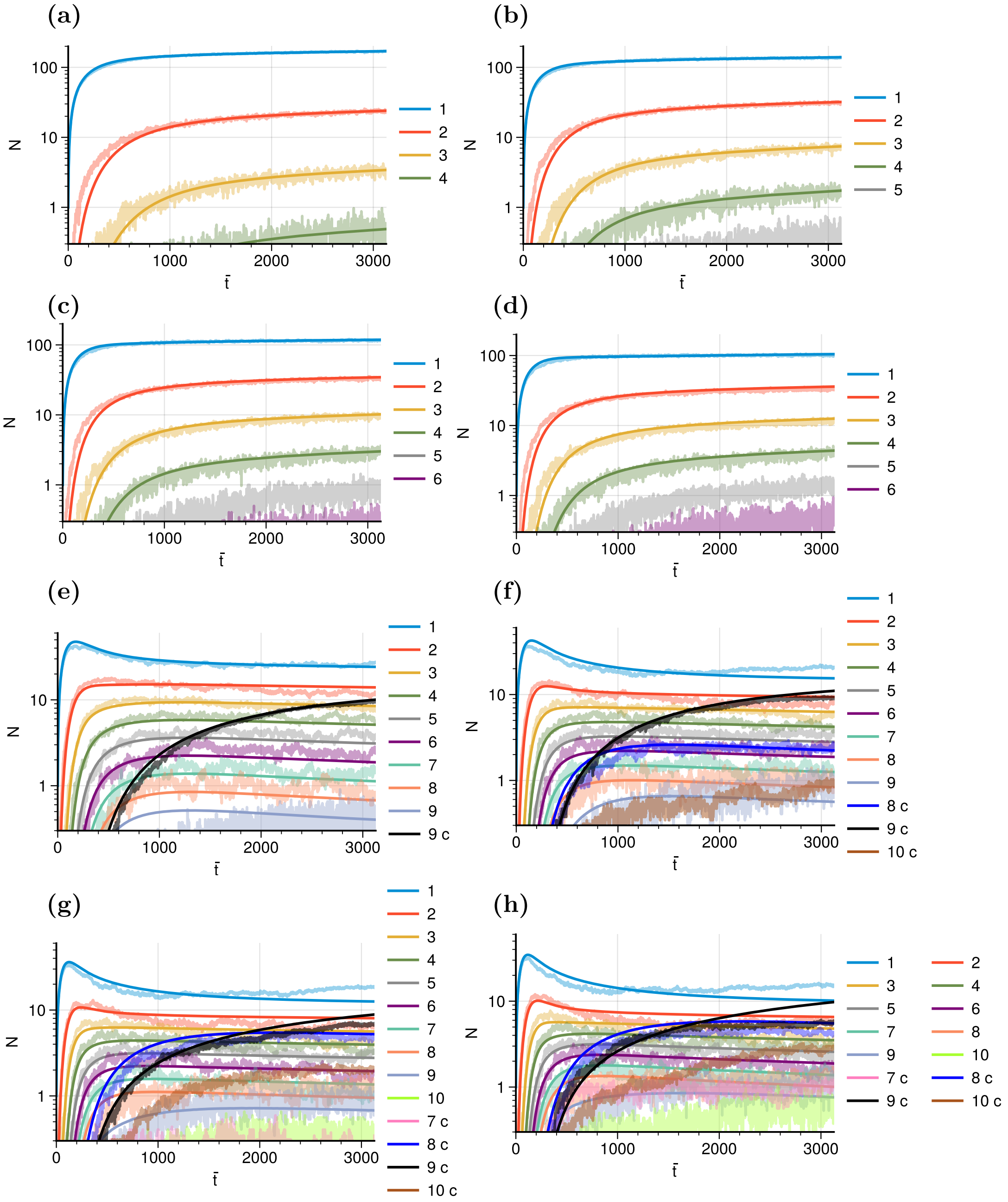


Figure S3: Simulated (light colored) and fitted (darker lines) concentrations for $L$ = 170nm as a function of rescaled time ($\bar{t} = t/16\text{ns}$). The parameters were $\epsilon_a/\epsilon_r = 1$ and (a) $\theta_c = 0.1\pi$,(b) $\theta_c = 0.2\pi$, (c) $\theta_c = 0.3\pi$, (d) $\theta_c = 0.2\pi$; $\epsilon_a/\epsilon_r = 2$ and (e) $\theta_c = 0.1\pi$,(f) $\theta_c = 0.2\pi$, (g) $\theta_c = 0.3\pi$, (h) $\theta_c = 0.2\pi$ The label "c" indicates a closed species.

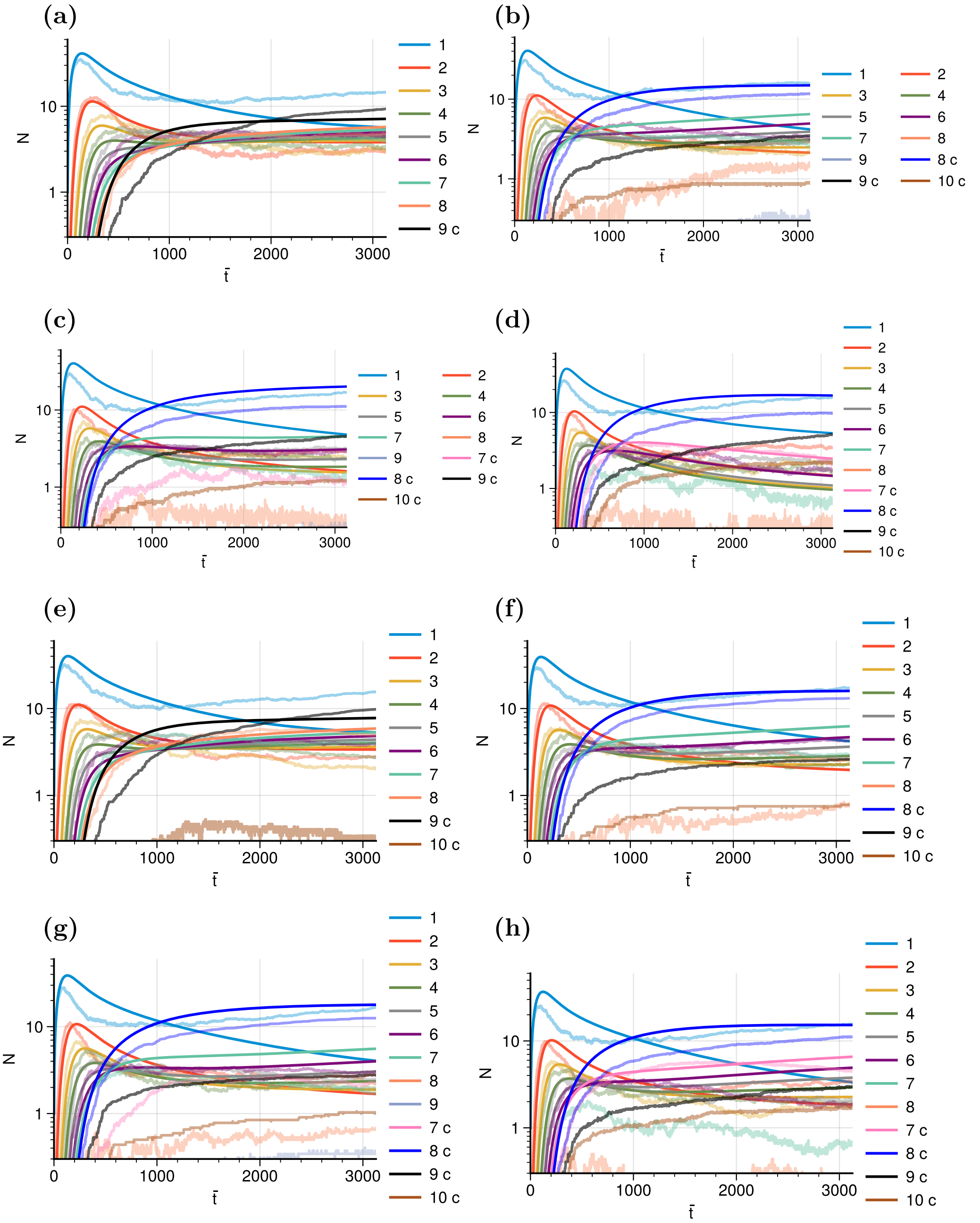


Figure S4: Simulated (light colored) and fitted (darker lines) concentrations for $L = 170\text{nm}$ as a function of rescaled time ($\bar{t} = t/16\text{ns}$). The parameters were $\epsilon_a/\epsilon_r = 3$ and (a) $\theta_c = 0.1\pi$,(b) $\theta_c = 0.2\pi$, (c) $\theta_c = 0.3\pi$, (d) $\theta_c = 0.2\pi$; $\epsilon_a/\epsilon_r = 4$ and (e) $\theta_c = 0.1\pi$,(f) $\theta_c = 0.2\pi$, (g) $\theta_c = 0.3\pi$, (h) $\theta_c = 0.2\pi$ The label "c" indicates a closed species.

| $\epsilon_a/\epsilon_r$ | $\theta_c/\pi$ 0.1 | 0.2 | 0.3 | 0.4 |
|---|---|---|---|---|
| 1.0 | 0.0272 | 0.0613 | 0.0967 | 0.119 |
| 2.0 | 0.639 | 0.815 | 1.22 | 1.34 |
| 3.0 | 0.825 | 0.866 | 0.874 | 1.04 |
| 4.0 | 0.885 | 0.935 | 0.965 | 1.11 |

(a) $k_+(\times 10^{-4}/16\text{ns})$

| $\epsilon_a/\epsilon_r$ | $\theta_c/\pi$ 0.1 | 0.2 | 0.3 | 0.4 |
|---|---|---|---|---|
| 1.0 | 31.7 | 36.0 | 38.2 | 35.0 |
| 2.0 | 23.9 | 16.7 | 19.6 | 16.0 |
| 3.0 | 2.62 | 0.856 | 0.302 | 0.0607 |
| 4.0 | 2.33 | 0.775 | 0.546 | 0.926 |

(b) $k_-(\times 10^{-4}/16\text{ns})$

Table S2: Optimized association and dissociation constants for all the combinations of microscopic parameters $\epsilon_a/\epsilon_r$ and $\theta_c$. All values have dimensions of inverse time.

| | Open rate $k_o$/16ns | | Close rate $k_c$/16ns | |
|---|---|---|---|---|
| $\theta_c/\pi$ | Octamer | Nonamer | Octamer | Nonamer |
| 0.1 | - | $4.9\times10^{-4}$ | - | 0.0489 |
| 0.2 | 2.343554 | $9.51\times10^{-4}$ | 9.32 | 0.0522 |
| 0.3 | 0.00491 | $1.\ 79\times10^{-4}$ | 0.0374 | 0.0108 |
| 0.4 | 0.00268 | $4\times10^{-6}$ | 0.0188 | 0.00728 |

Table S3: Optimized opening and closing rates for $\epsilon_a/\epsilon_r = 2$, where reversibility was observed. All values have dimensions of inverse time. For $\theta_c = 0.1\pi$ only closed nonamers are observed.
